# V-SWITCH: A single-vector OFF-to-ON fluorescent reporter of live RNA virus infections

**DOI:** 10.64898/2026.04.08.717260

**Authors:** Miguel Cid-Rosas, Michelle Grunberg, Alejandro Matía, See-Chi Lee, Hunter Woosley, Soorya Pradeep, Shivali Kanwar, Sudip Khadka, Daniele Jones, Shalin Mehta, Ruth Hüttenhain, Vincent Turon-Lagot, Carolina Arias

**Author notes:** Department of Microbiology and Immunology, University of California, San Francisco, San Francisco, CA, USA. Department of NanoBiophotonics, Max Planck Institute for Multidisciplinary Sciences, Göttingen, Germany (Hunter Woosley).

## Abstract

Fluorescent reporters enable direct analysis of viral infection, but existing approaches often require viral genome engineering or lack sensitivity during authentic infection. Here, we present V-SWITCH, a modular, single-vector, host-encoded OFF-to-ON fluorescent reporter for live RNA virus infection. V-SWITCH uses a split mNeonGreen release-and-capture mechanism in which viral protease activity releases an endoplasmic-reticulum-anchored mNG3A(1–10) fragment, enabling complementation with a constitutively expressed nuclear mNG(11)-BFP module. The BFP signal controls for reporter expression and supports automated nuclear segmentation. We validated V-SWITCH for DENV, ZIKV, WNV, and HCoV-OC43 across multiple cell types. Reporter activation closely matches viral-antigen detection and the fluorescence intensity correlates with intracellular levels of viral RNA. The reporter enabled flow-cytometry-based quantification of infection efficiency, fluorescence-activated cell sorting of infected/uninfected cells, and longitudinal single-cell imaging of heterogeneous infection dynamics. V-SWITCH further supported a pooled CRISPRi screen targeting 2,583 druggable genes and evaluation of antiviral activity of selected compounds. Notably, pharmacological follow-up provided previously undescribed evidence that Plitidepsin, an EEF1A inhibitor, and GSK-A1, a PI4KA inhibitor, restrict DENV infection in cultured cells. V-SWITCH provides a scalable, robust, and versatile platform for studying protease-expressing RNA viruses and screening for virus- and host-directed antivirals.

## Introduction

A central challenge in virology is the development of reporter systems that can capture the dynamics of infection while being easy to deploy, are adaptable across experimental contexts, and cause minimal perturbations to the host. Fluorescence-based optical reporters enable real-time, non-destructive measurements of infection at both the population and single-cell level. However, the development of optical reporters of infection remains constrained by limited versatility, reliance on virus engineering, or incompatibility with longitudinal and high-throughput analyses. Thus, the need remains for reporters allowing the interrogation of infection across time in diverse viruses and cell types.

Flaviviruses are an example of both the urgency of this need and the challenges inherent in monitoring viral infection and replication. Viruses from this family, Dengue virus (DENV), Zika virus (ZIKV), West Nile virus (WNV), Yellow Fever virus (YFV), and Japanese encephalitis virus (JEV), among others, represent a major global public health threat, demonstrate sustained endemic transmission, and lack approved antiviral therapies ^1,2^. To date, a wide range of optical reporters have been generated by directly modifying flavivirus genomes to encode fluorescent proteins, including GFP-labeled DENV, ZIKV, JEV, and YFV, as well as mCherry-labeled DENV ^3–7^. These tools allow the dynamic study of viral replication and spread. However, their reliance on the insertion of large heterologous sequences into compact viral genomes frequently alter replication kinetics or is genetically unstable, thus limiting their applicability, particularly for comparative analyses, time courses, or large-scale studies ^8^.

To circumvent the limitations associated with viral-genome engineering, host-encoded reporters have emerged as a compelling alternative. These “switch-based” biosensors exploit virus-specific protease activity to trigger a detectable signal while preserving the native viral genome ^9^. Arias-Arias and colleagues developed FlaviA, in which protease-mediated removal of a quencher activates fluorescence during infection with unmodified DENV, ZIKV or YFV ^10^. FlaviA enabled imaging-based detection of infection, and a far-red mNeptune variant improved plaque visualization. These advances established the potential of protease-activated fluorescence for monitoring authentic flavivirus infection, although flow-cytometric quantification and isolation of infected cells were not evaluated. Translocation-based reporters have also enabled reliable live-virus detection ^11,12^, but measure redistribution of a constitutive fluorescent signal rather than an increase in total fluorescence, complicating direct population-based selection. FlipGFP-based systems have provided high-throughput protease assays but were primarily validated using protease overexpression or non-replicating surrogates ^13,14^. A related reporter was recently adapted to diverse picornaviral 3C proteases, although activation during EV-A71 infection was detected in only a subset of infected cells^15^. Building on these approaches, a host-encoded reporter that combines robust fluorescence activation during authentic infection with quantitative flow cytometry and FACS-based isolation would enable reporter-based pooled genetic screening.

Here, we present V-SWITCH: a sensitive, single-vector, cell-encoded split-fluorescent reporter system that detects viral protease activity during authentic viral infection. Using DENV as one of our primary model systems, we combine the specificity of enzyme-mediated cleavage with the high signal-to-noise properties of split-protein complementation. At a molecular level, as is the case with all flaviviruses, DENV replication depends on the processing of a single viral polyprotein by host signal peptidases and the viral NS2B/3 serine protease ^16^. The NS2B/3 protease cleaves the polyprotein at multiple sites (e.g., NS2A/NS2B, NS2B/NS3, NS3/NS4A, NS4B/NS5), making its activity indispensable for viral maturation and replication ^17^. Protease activity is tightly coupled to productive infection, thus representing an ideal proxy for monitoring viral replication kinetics and evaluating the efficacy of potential inhibitors ^18^.

Building on the split-mNeonGreen (mNG) system ^19^, we engineered V-SWITCH around a “release-and-capture” mechanism in which a detector fragment anchored to the endoplasmic reticulum (ER) is liberated by viral protease activity and subsequently localized to the nucleus, generating a fluorescent signal. Unlike prior sensors restricted to protease overexpression, this platform robustly detects protease activity during live virus infection. The resulting high contrast OFF-to-ON fluorescence switch enables rapid, antibody-free quantification of infection by flow cytometry and allows for physical separation of infected and uninfected cells by fluorescence-activated cell sorting (FACS), facilitating downstream applications such as pooled CRISPR screens and transcriptomic profiling. Finally, we demonstrate the modularity of V-SWITCH by adapting it to detect ZIKV, WNV, and the human coronavirus OC43. This single-vector reporter is modular and easily adaptable, facilitates resource sharing, and establishes a broadly applicable, rapidly deployable tool for studying viral protease activity and reporting on infection dynamics across diverse viruses and cell types.

## Results

### Development of DENV-SWITCH as a modular, single-vector reporter platform for Dengue infection

To monitor DENV infection in real-time with single-cell resolution, we engineered DENV-SWITCH, a reporter system based on the specific cleavage of an ER-anchored fluorescent sensor by the DENV NS2B/3 protease, similar to previously described fluorescence-translocation reporters ^11,12^. DENV-SWITCH relies on a split mNG expressed in two different modules within a single vector (Fig. 1A) ^19^. The Cleavable Anchor Module (CAM), is composed of the long mNG3A(1-10) fragment fused at its N-terminus to a Nuclear Localization Signal (NLS) followed by a DENV protease cleavage site (PCS), the transmembrane (TM) domain of Sec61β as an ER-anchor, a T2A sequence, and a Blasticidine resistance gene for selection. The Nuclear Capture Module (NCM) consists of an mNG(11) fragment linked to a Nuclear Localization Signal (NLS) and a TagBFP, which increases mNG(11) stability, serves as a constitutive marker of reporter expression, and facilitates nuclei segmentation for high-content image analysis (Fig. 1A). In this single-vector architecture, the CAM and the NCM are expressed from a lentiviral construct driven by the EF1α and CMV promoters, respectively (Fig. 1A-B). To improve fluorescence reconstitution, we used mNG3A(1-10), which has been reported to have a higher affinity for mNG(11) and thus enhances complementation ^20^. Alternative versions of this system, one using a mNG2(1-10) as the NCM paired with the shorter mNG(11) as the cleavable ER-anchored element (CAM), and a second using mNG(11) tagged to a nuclear or nucleolar protein, successfully detected infection in multiple contexts; however both showed limited stability and sensitivity, indicating that DENV-SWITCH represents an optimal version of this reporter (Sup Fig. 1 and 2).

**Figure 1.**
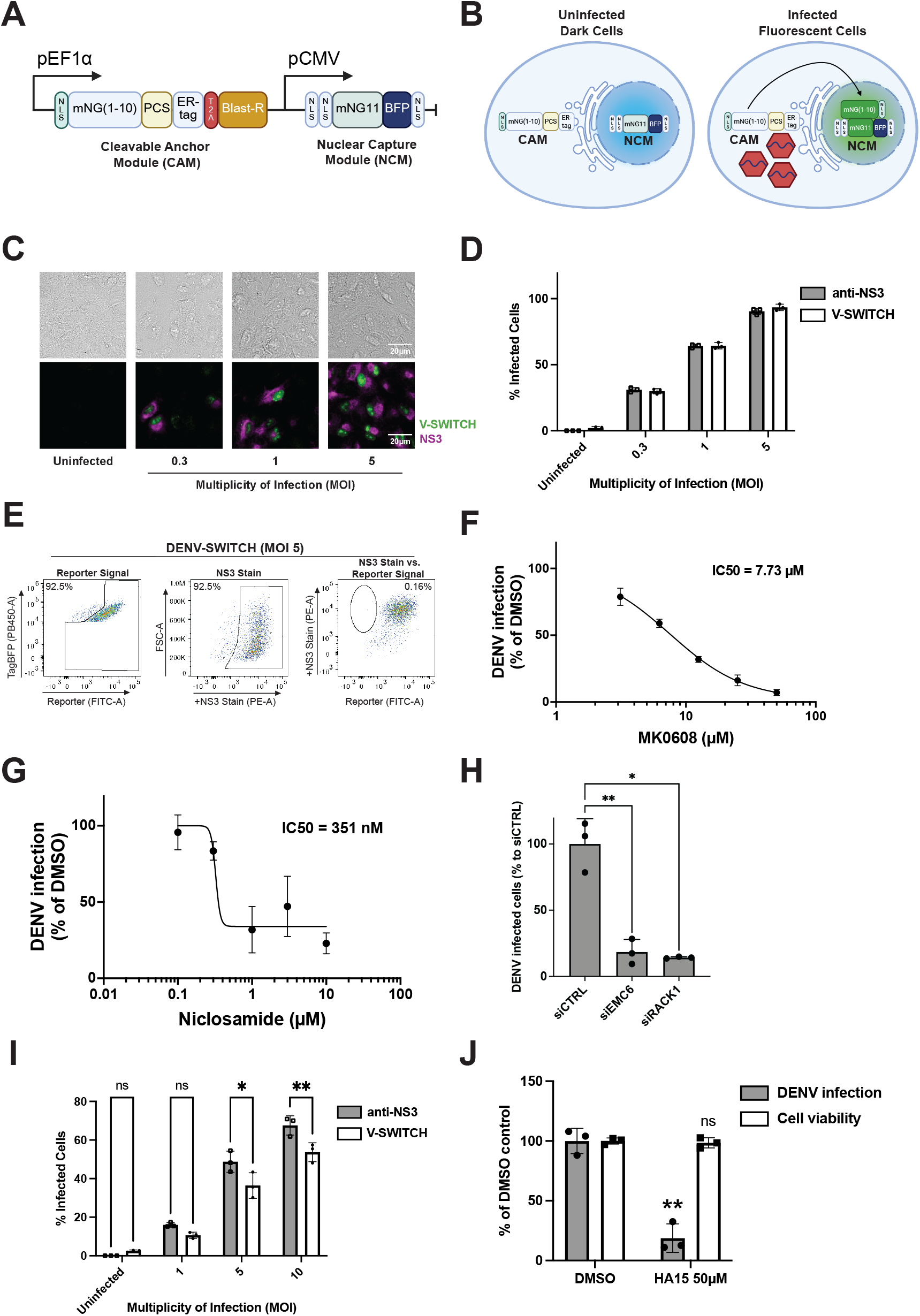
DENV-SWITCH: a modular, single-vector reporter platform for flaviviruses. (A–B) Architecture and activation of DENV-SWITCH. A single lentiviral vector encodes the Cleavable Anchor Module (CAM), containing ER-anchored mNG3A(1–10) and a viral protease cleavage site, and the Nuclear Capture Module (NCM), containing nuclear mNG(11)–BFP. Protease cleavage releases mNG3A(1–10), enabling nuclear fluorescence complementation. (C) Representative images of A549 DENV-SWITCH cells infected with DENV at the indicated MOIs. Reporter fluorescence is green and NS3 immunostaining magenta. Scale bar, 20 µm. (D) Infection detected by NS3 immunostaining (grey) or reporter fluorescence (white). (E) Representative flow cytometry plots at MOI 5, showing reporter fluorescence, NS3 immunostaining and their relationship. BFP-positive cells define the reporter-expressing population. (F–G) MK0608 and niclosamide dose–response curves in A549 DENV-SWITCH cells. Absolute IC50 values were 7.73 µM (95% profile-likelihood CI, 6.99–8.52 µM) and approximately 351 nM, respectively; the niclosamide confidence interval could not be determined. Absolute IC50 is the fitted concentration giving 50% of the DMSO-control response. (H) Effects of siRNA-mediated EMC6 or RACK1 silencing on DENV infection at 24 hpi (MOI 5), expressed relative to mean siCTRL infection. (I) Infection detected by NS3 immunostaining (grey) or reporter fluorescence (white) in BJ-5ta fibroblasts at the indicated MOIs. (J) Effects of the BiP/GRP78 inhibitor HA15 (50 µM) on DENV infection (black; relative to DMSO) and cell viability (white) in BJ-5ta cells at 24 hpi. Quantitative data are mean ± s.d. from three independent experiments. Statistical comparisons used two-way repeated-measures ANOVA with matching in both factors and Geisser–Greenhouse correction, followed by Šídák-adjusted comparisons between detection methods within each MOI (D, I); Geisser–Greenhouse-corrected repeated-measures one-way ANOVA with Dunnett-adjusted comparisons against siCTRL (H); and paired two-tailed t-tests for infection and viability (J). Exact P values are provided in Source Data.

**Figure 2.**
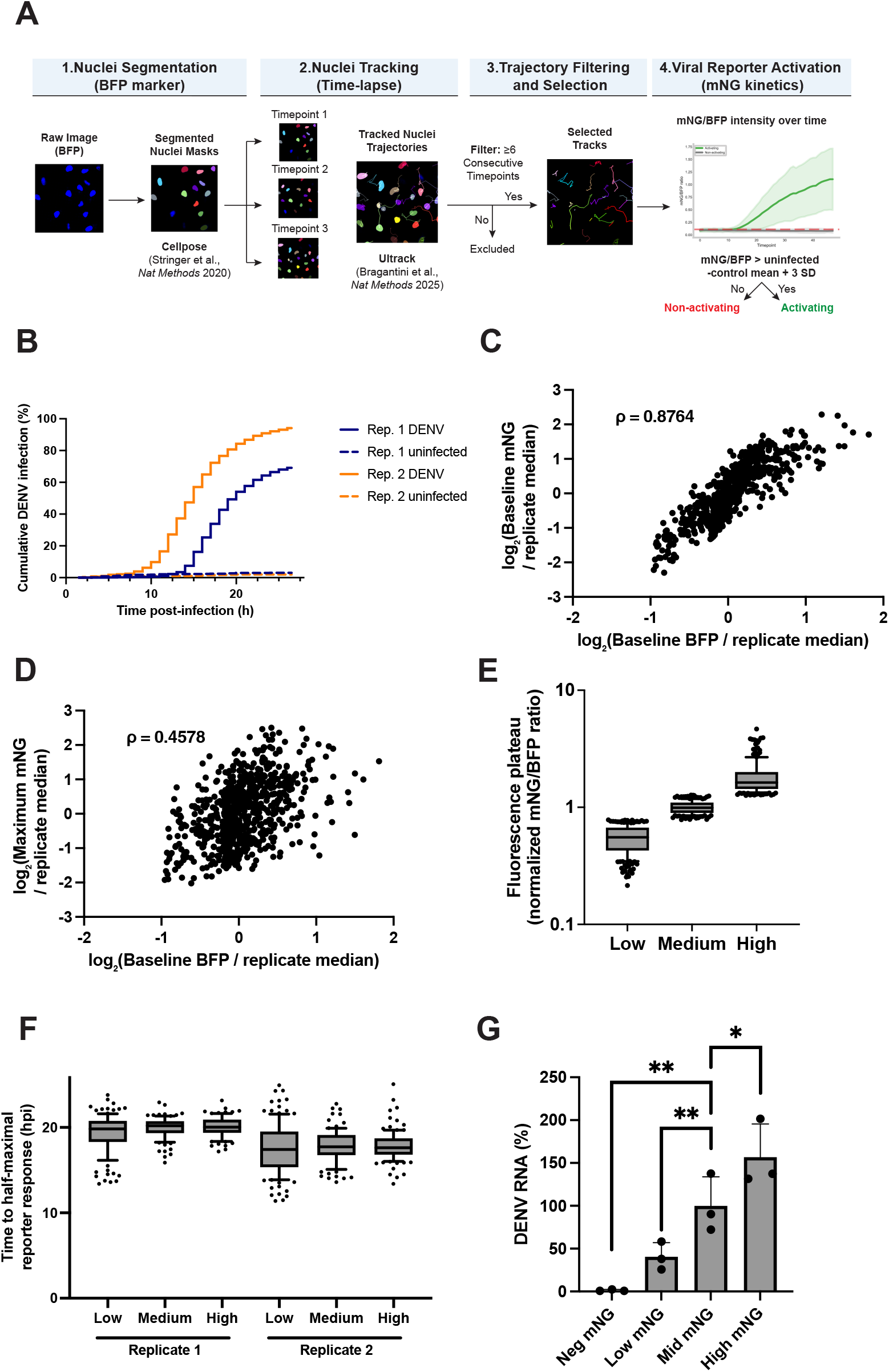
Live-cell imaging reveals heterogeneous reporter-response dynamics and reporter intensity is associated with intracellular DENV RNA. (A) Image-analysis workflow for two independent experiments (27 hourly frames; 1.5–27.5 hpi). BFP-positive nuclei were segmented and tracked. Activation required two consecutive mNG/BFP measurements above the matched uninfected-control mean plus three standard deviations. (B) Cumulative reporter activation in DENV-exposed cells (MOI 5; solid lines) and uninfected controls (dashed lines), by replicate. Kaplan–Meier estimates account for different tracking periods. Cells already reporter-positive when tracking began were excluded. (C–D) Baseline BFP versus baseline mNG (C) or maximum observed mNG (D), using 724 cells (392 and 332 by replicate). Baseline intensities were averaged over frames 0–5. Intensities were normalized to within-replicate medians and log2-transformed. Spearman’s ρ = 0.8764 (C) and 0.4578 (D). (E–F) Fitted fluorescence plateaus (E) and times to half-maximal reporter response, t50 (F). Low, Medium and High groups were defined by plateau tertiles among initially eligible cells within each replicate. The same 626 cells with reliable t50 estimates are shown in both panels, retaining their original groups: 287 cells in replicate 1 (115/92/80 by group) and 339 in replicate 2 (121/99/119). Plateaus were normalized within each replicate to the initial cohort’s median and pooled in E; F shows t50 by replicate. Boxes show medians and interquartile ranges; whiskers mark the 10th–90th percentiles, with values beyond them shown individually. This exploratory subset represents 21.4% of initially eligible cells; t50 measures reporter-response timing, not infection onset. Panels B–F are descriptive, without cell-level significance tests. (G) DENV RNA in populations sorted by mNG/BFP ratio, normalized to GAPDH and expressed relative to the mid-mNG population. Data are mean ± s.d. from three independent experiments. Paired ΔCq values were analyzed by Geisser–Greenhouse-corrected repeated-measures one-way ANOVA with Dunnett-adjusted comparisons against mid-mNG. Exact P values and track counts are provided in Source Data.

We established a stable DENV-SWITCH reporter A549 cell line by transduction with a lentiviral vector and single-cell clone selection. We validated DENV-SWITCH performance upon DENV infection, confirmed by NS3 immunofluorescence. The nuclear accumulation of mNG fluorescence was observed almost exclusively in cells positive for viral NS3, with high spatial concordance and minimal background in uninfected cells (Fig. 1C). We quantified the sensitivity of the system by flow cytometry across a range of multiplicity of infection (MOI) and observed a strong agreement between the percentage of infected cells detected by NS3 immunostaining and reporter fluorescence, with concordance ranging from 93.5 to 99.7% (Fig. 1D). These results confirm that DENV-SWITCH accurately detects viral infection. The constitutive BFP expression improves quantification of infected cells by excluding cells with improper reporter expression from the assay (Fig. 1E ; Sup Fig. 3).

**Figure 3.**
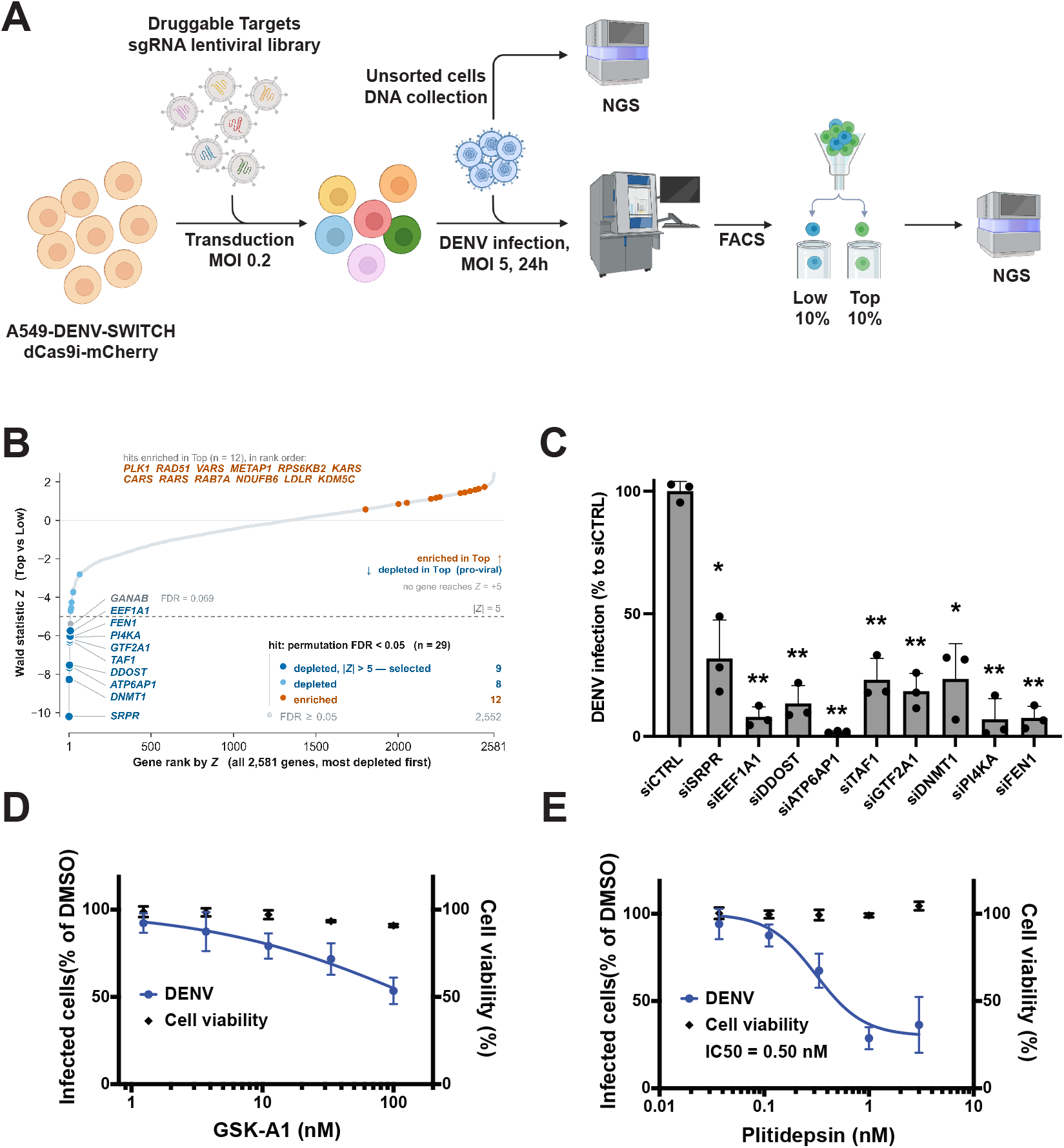
DENV-SWITCH enables pooled CRISPRi screening for druggable host dependencies of DENV infection. (A) Pooled CRISPRi screen in A549 DENV-SWITCH cells expressing dCas9i–mCherry. A library of 14,545 sgRNAs targeting 2,583 druggable genes was screened following DENV infection (MOI 5). The 10% lowest (Low) and highest (Top) reporter-activation populations were sorted in two biological replicates, and sgRNA abundance was quantified by sequencing. (B) Ranked gene-level Wald Z statistics from the MAGeCK-MLE Top-versus-Low comparison. Hits met a permutation FDR ≤ 0.05. Negative and positive scores indicate candidate dependency and restriction factors, respectively. The nine negative hits selected for follow-up are indicated. (C) Effects of siRNA-mediated silencing of SRPR, EEF1A1, DDOST, ATP6AP1, TAF1, GTF2A1, DNMT1, PI4KA or FEN1 on DENV infection at 24 hpi (MOI 5), expressed relative to mean siCTRL infection. Geisser–Greenhouse-corrected repeated-measures one-way ANOVA was followed by Dunnett-adjusted comparisons against siCTRL; exact P values are provided in Source Data. (D–E) Dose–response analysis of GSK-A1, a PI4KA inhibitor (D), and plitidepsin, an EEF1A-targeting compound (E), in DENV-infected BJ-5ta DENV-SWITCH cells. At 100 nM GSK-A1, infection was 53.6 ± 7.6% of the DMSO control and viability was 90.9 ± 0.8%. Plitidepsin had an absolute IC50 of 0.504 nM (95% profile-likelihood CI, 0.347–0.822 nM), defined as the fitted concentration giving 50% of the DMSO-control response. At 1 nM, infection was 28.7 ± 6.3% of control without a detectable reduction in viability. Data in C–E are mean ± s.d. from three independent experiments. Technical replicates were averaged within each experiment before fitting infection-response curves. Viability is shown as measured points without a fitted curve; CC50 values were not estimated because viability did not approach 50%.

We then evaluated the suitability of DENV-SWITCH for antiviral compound testing. Treatment with MK0608 and niclosamide reduced reporter activation, consistent with their reported antiviral activity (Fig. 1F–G) ^21,22^. Absolute IC50, defined as the fitted concentration giving 50% of the DMSO-control response, was 7.73 µM for MK0608 and 351 nM for niclosamide. To test our system for the identification of viral dependencies using gene-expression inhibition, we silenced EMC6 and RACK1, two host factors previously implicated in DENV replication ^23,24^. The siRNA-mediated knockdown of these factors resulted in a significant reduction of DENV-SWITCH activation by 81.6% and 85.8%, respectively, at 24 hours post infection (Fig. 1H, Sup Fig 4)

**Figure 4.**
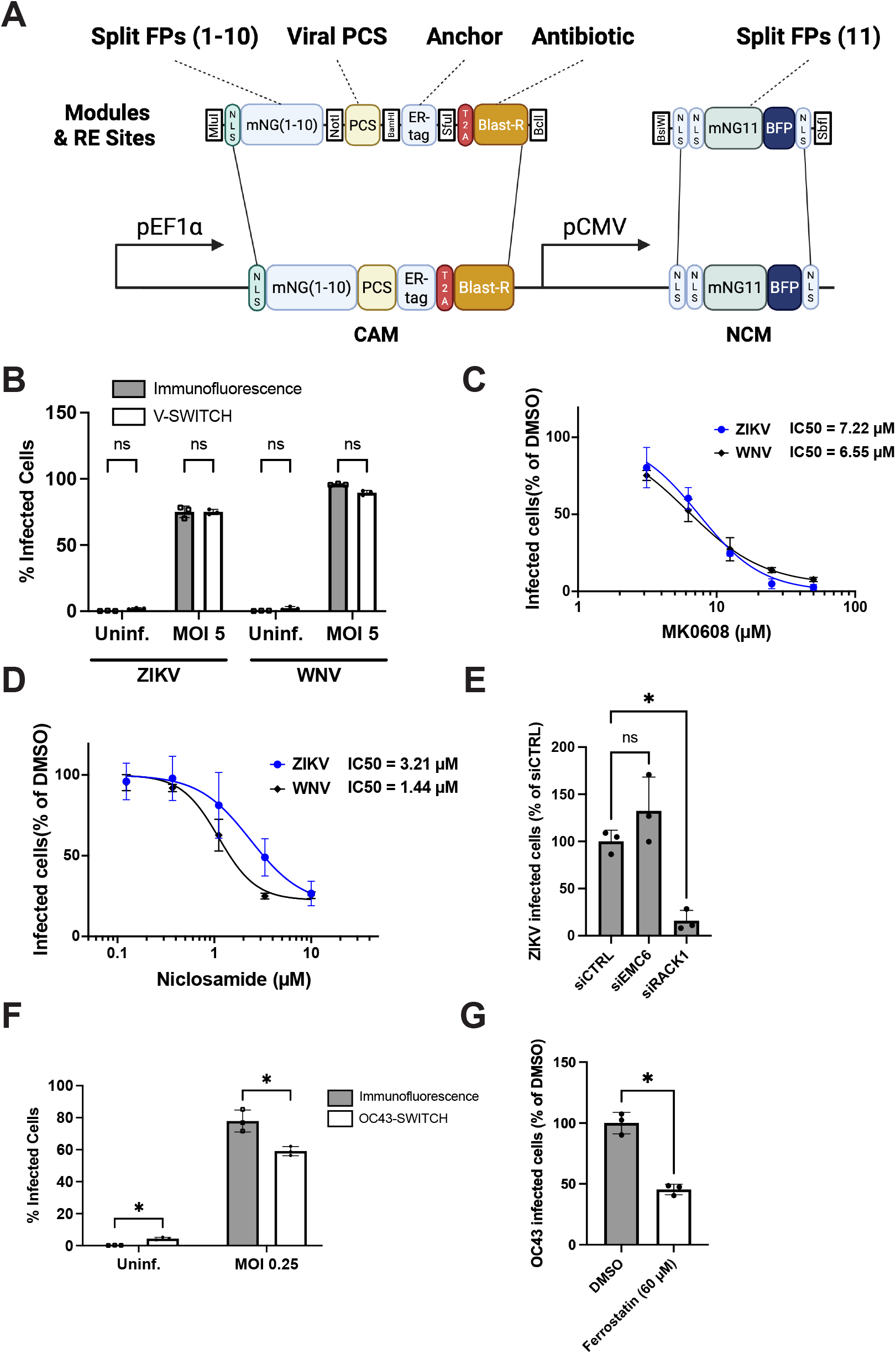
Adaptability of the V-SWITCH platform to other viral proteases. (A) Modular V-SWITCH architecture. Unique restriction sites flank the split-fluorescent-protein fragments, viral protease cleavage site, membrane anchor and antibiotic resistance cassette, enabling cassette exchange. CAM and NCM gene maps are shown below. (B) Infection detected by immunostaining (grey; anti-NS2B for ZIKV and anti-E for WNV) or reporter fluorescence (white) in A549 ZIKV-SWITCH and WNV-SWITCH cells at MOI 5. (C–D) MK0608 (C) and niclosamide (D) dose–response curves in these cell lines. MK0608 absolute IC50 values were 7.22 µM (95% profile-likelihood CI, 6.36–8.17 µM) for ZIKV and 6.55 µM (5.88–7.31 µM) for WNV. Niclosamide values were 3.21 µM (1.98–5.10 µM) and 1.44 µM (1.21–1.79 µM), respectively. Absolute IC50 is the fitted concentration giving 50% of the DMSO-control response. (E) Effects of siRNA-mediated EMC6 or RACK1 silencing on ZIKV infection in A549 ZIKV-SWITCH cells, expressed relative to mean siCTRL infection. (F) Infection detected by anti-HCoV-OC43 immunostaining (grey) or reporter fluorescence (white) in HEK 293T OC43-SWITCH cells at 48 hpi (MOI 0.25). (G) Effects of ferrostatin-1 (60 µM) on OC43 infection, expressed relative to mean DMSO-control infection. Data are mean ± s.d. from three independent experiments. Detection methods were compared using paired two-tailed t-tests with Holm–Šídák adjustment across four virus-by-infection conditions (B) or two infection conditions (F). In E, Geisser–Greenhouse-corrected repeated-measures one-way ANOVA was followed by Dunnett-adjusted comparisons against siCTRL (adjusted P = 0.2886 for siEMC6 and 0.0150 for siRACK1). Ferrostatin-1 was compared with DMSO using a paired two-tailed t-test (G; P = 0.0159). Remaining exact P values are provided in Source Data.

To further assess the versatility of this reporter, we evaluated its performance for antiviral testing in a non-cancerous cell line. Virus-host interaction studies have uncovered new viral dependencies that could be used for therapeutic purposes. Many host-targeting compounds have been previously identified as anticancer agents and can be toxic to transformed cell lines routinely used for viral studies ^25^. To circumvent this limitation, we introduced the DENV-SWITCH reporter into BJ-5ta, an immortalized human dermal fibroblast cell line. BJ-5ta cells were infected with DENV (MOI 1, 5, and 10), and infection was evaluated by DENV-SWITCH activation and DENV-NS3 immunofluorescence. The reporter detected infected cells across all MOIs, though with less sensitivity than with NS3 staining. This discrepancy could reflect a lower baseline of reporter expression in this cell line, which does not affect NS3 staining (Fig. 1I).

To test the utility of this reporter for evaluating the activity of host-directed antivirals, we treated BJ-5ta-DENV-SWITCH cells with HA15, a BiP/GRP78 ATPase domain inhibitor, with broad-spectrum antiviral activity against ZIKV and multiple dsDNA viruses ^26–28^. BiP/GRP78 is a chaperone localized at the endoplasmic reticulum, upregulated during DENV infection ^29^. At a concentration of 50 µM, HA15 was effective against DENV, reducing DENV-SWITCH activation by 70% without a detectable reduction in cell viability (Fig. 1J). These results demonstrate the antiviral activity of HA15 against DENV and validate the use of this reporter in a non-cancerous cell line for evaluating host-targeting compounds with broad-spectrum antiviral potential.

Collectively, these data demonstrate that the single-vector DENV-SWITCH is a robust tool capable of quantifying live DENV infection and suitable for studying host-pathogen interactions and antiviral activity of host-and viral-targeted agents.

### Live-cell imaging reveals heterogeneity in reporter-response dynamics

A distinct advantage of this fluorescence-based reporter is the ability to monitor the temporal dynamics of infection in live cells at single-cell resolution. We infected A549 DENV-SWITCH cells with DENV at MOI 5 and analyzed two independent biological experiments collecting images every hour over 27 timepoints, corresponding to 1.5–27.5 hours post-infection (hpi). In our analysis workflow, we performed nuclei segmentation using Cellpose ^30^ on the constitutively expressed BFP nuclear marker. Individual nuclei were tracked using Ultrack ^31^. Tracks with at least six consecutive observations were retained. Reporter activation was detected when the mNG/BFP ratio exceeded the matched uninfected-control mean plus three standard deviations for two consecutive frames (Fig. 2A).

Single-cell tracking revealed heterogeneity in reporter-detection kinetics in both biological replicates. Cumulative activation was estimated using left-truncated Kaplan–Meier curves, allowing for staggered track entry and right censoring. At 26.5 hpi, the estimated cumulative activation was 69.1% and 94.1% in the two DENV-infected replicates, respectively, compared with 3.1% and 1.9% in their matched uninfected controls (Fig. 2B). These values describe reporter detection rather than the time of infection initiation. To examine the contribution of basal reporter abundance, we analyzed 724 cells with a complete preactivation baseline (392 and 332 cells in replicates 1 and 2). After normalizing each intensity measure to its within-replicate median, baseline BFP and baseline mNG were strongly associated (Spearman’s ρ = 0.8764; Fig. 2C). However, baseline BFP showed a more moderate association with the observed maximum mNG intensity during tracking (ρ = 0.4578) pointing to a fluorescence activation dynamic independent from basal reporter expression (Fig. 2D).

We next examined whether differences in reporter intensity were associated with differences in response timing. An earlier response could allow more time for fluorescent signal to accumulate, potentially contributing to variations between cells. We therefore grouped cells into Low, Medium and High responders according to the fluorescence plateau estimated from their individual trajectories (Fig. 2E), and compared when each cell reached half of its fluorescence increase above baseline (t50). This comparison uses the same stage of each cell’s response, rather than requiring weak and strong responders to reach the same fluorescence intensity. Among the 626 cells whose response timing could be reliably estimated, the three groups reached this halfway point at similar median times: 19.8–20.2 hpi in replicate 1 and 17.4–17.7 hpi in replicate 2 (Fig. 2F). Thus, markedly different response intensities were observed in groups with similar reporter-response timing and could reflect differences in viral replication levels.

To test whether differences in reporter intensity corresponded to differences in intracellular viral RNA, we sorted DENV-infected cells at 24 hpi into mNG-negative, low-, medium- and high-fluorescence populations based on their mNG/BFP ratio (Sup. Fig. 5). RT–qPCR detected no DENV RNA in the mNG-negative population. Surprisingly, and compared with the medium-fluorescence population, the low-fluorescence population contained 60% less DENV RNA, whereas the high-fluorescence population contained 58% more (Fig. 2G). These results establish that reporter mean fluorescence intensity can distinguish populations with different intracellular DENV RNA levels, supporting its use to isolate subpopulations of cells for downstream studies of host factors affecting infection.

### DENV-SWITCH enables pooled CRISPRi screening for DENV host dependencies

Given that DENV-SWITCH fluorescence intensity correlated with intracellular DENV RNA abundance, we evaluated whether the reporter is suited for pooled genetic screening and identification of host factors that regulate viral replication. We generated A549 DENV-SWITCH cells expressing dCas9i–mCherry and transduced them with a pooled sgRNA library targeting 2,583 druggable targets. Following DENV infection at high MOI (MOI = 5), we sorted BFP-high cells into two populations: the 10% with the lowest V-SWITCH activation (Low) and the 10% with the highest (Top). Unlike a survival screen, which identifies host factors based on survival or proliferation, this experimental design selects cells based directly on reporter activation, isolating sgRNAs enriched in cells that restrict (low) or promote (high) viral replication. The screen was performed in two independent biological replicates, and sgRNA abundance in each population was quantified by next generation sequencing (NGS) (Fig. 3A).

The sgRNA counts were analyzed using MAGeCK-MLE, comparing sgRNA enrichment between the Low and the Top sorted populations. The analysis identified 29 genes with an FDR below 0.05, including 17 genes with negative Z-scores and 12 with positive Z-scores (Fig. 3B). Negative Z-scores indicated depletion from the Top population, meaning potential pro-viral host factors, whereas positive scores indicated enrichment in the Top population, consistent with candidate antiviral restriction factors. To prioritize candidates for experimental follow-up, we applied a threshold of |Z| > 5. Nine genes, SRPR, DNMT1, ATP6AP1, DDOST, TAF1, GTF2A1, PI4KA, FEN1, and EEF1A1, met both criteria and all displayed negative Z-scores.

The prioritized candidates spanned several cellular processes, including ER protein targeting and N-linked glycosylation (SRPR and DDOST), V-ATPase-dependent organelle acidification (ATP6AP1), translation elongation (EEF1A1), transcriptional and epigenetic regulation (TAF1, GTF2A1, and DNMT1), phosphoinositide metabolism (PI4KA), and DNA replication and repair (FEN1) ^32–38^. DDOST and ATP6AP1 have previously been identified in genetic screens for DENV host factors, whereas SRPR and EEF1A1 have been implicated in DENV or flavivirus infection through protein-interaction or functional studies ^39–42^. Earlier DENV reporter-virus and replicon studies in other cellular contexts, tested PI4KA and found little or no requirement for this kinase ^43,44^. By contrast, TAF1, GTF2A1, DNMT1, and FEN1 have never been identified as DENV host-dependency factors in human cells.

We evaluated the nine potential host factors by siRNA-mediated silencing in A549 DENV-SWITCH cells. Silencing of each gene reduced DENV infection by 75–99% relative to the non-targeting control (Fig. 3C). Knockdown efficiency was confirmed by RT-qPCR (Sup Fig. 6A). To assess whether gene silencing had an impact on the reporter expression, we quantified BFP fluorescence (Sup Fig. 6B). SRPR silencing reduced mean BFP intensity by approximately 60%, indicating that its effect on V-SWITCH activation could be due to reduced reporter abundance. In contrast, GTF2A1 silencing increased mean BFP intensity by approximately 100% (P = 0.0339). The approximately 40% higher mean after DNMT1 silencing was not statistically significant (P = 0.1852). DENV infection was measured as the absolute percentage of mNG-positive cells within the BFP-positive population, so these increases do not explain the reduction in DENV-SWITCH positive cells. The remaining siRNAs did not significantly change mean BFP intensity. Together, these results provide independent functional validation for eight of the nine candidates prioritized by the CRISPRi screen.

To determine whether the identified host factors could be targeted pharmacologically as antiviral targets, we evaluated inhibitors of PI4KA and EEF1A in BJ-5ta DENV-SWITCH fibroblasts. At 100 nM, GSK-A1 reduced DENV infection by 46.4%, while cell viability remained at 90.9 ± 0.8% of the DMSO control (Fig. 3D). An absolute IC50 was not reported because the mean reporter-defined infection did not reach 50% of the DMSO control within the tested concentration range. Plitidepsin inhibited DENV infection in a dose-dependent manner, with an absolute IC50 of 0.504 nM and a maximal observed reduction of 71.3% at 1 nM, without a detectable reduction in cell viability (Fig. 3E) Together, these findings demonstrate that V-SWITCH enables pooled functional-genomic screening and can facilitate the identification of host-targeted antiviral targets.

### Establishing V-SWITCH as an adaptable and versatile reporter of viral protease activity

A core objective of our modular design for this reporter was to ensure adaptability to multiple viral proteases. Unique restriction sites flanking each module allow the separation of the split fluorescent protein (splitFP), protease cleavage site (PCS), membrane anchor, and antibiotic resistance cassette coding regions, to be swapped independently through straightforward cloning strategies (Fig. 4A). To demonstrate the versatility of this platform, we adapted V-SWITCH to detect other viral pathogens by substituting the DENV PCS with those recognized by the Zika virus (ZIKV) NS2B/3 protease ^10^, West Nile virus (WNV) NS2B/3 protease ^45^, or the seasonal coronavirus OC43 nsp5/3CL protease (see Materials and Methods) (Table 1).

**Table 1.**
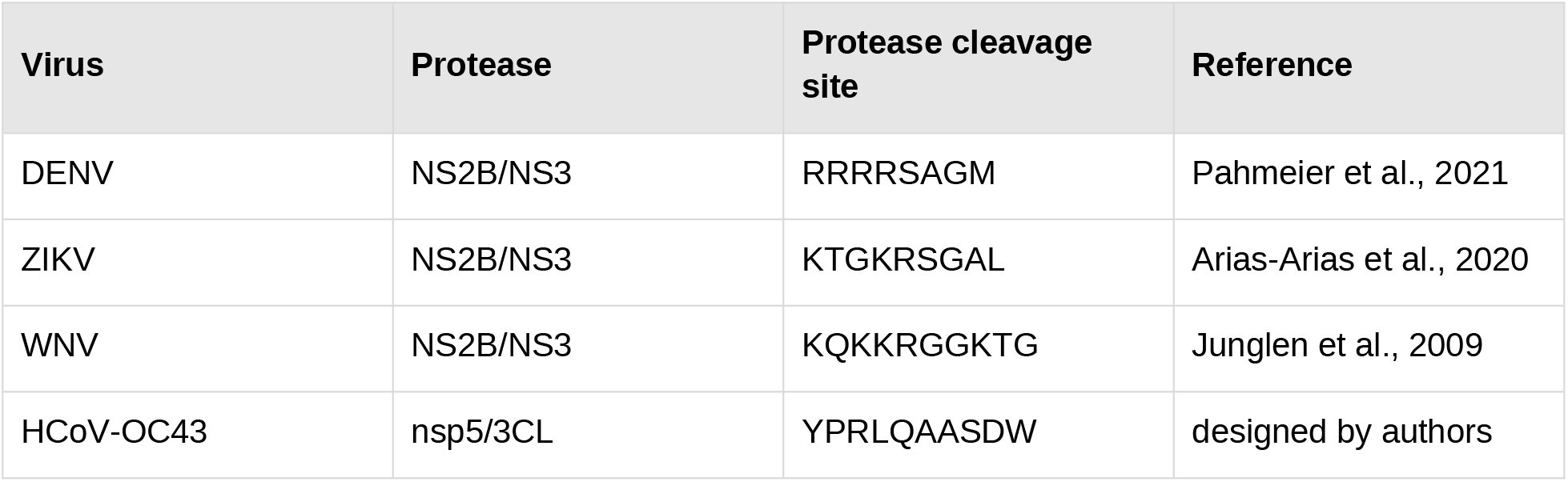
Viral protease cleavage sites used in V-SWITCH.

| **Virus** | **Protease** | **Protease cleavage site** | **Reference** |
|---|---|---|---|
| DENV | NS2B/NS3 | RRRRSAGM | Pahmeier et al., 2021 |
| ZIKV | NS2B/NS3 | KTGKRSGAL | Arias-Arias et al., 2020 |
| WNV | NS2B/NS3 | KQKKRGGKTG | Junglen et al., 2009 |
| HCoV-OC43 | nsp5/3CL | YPRLQAASDW | designed by authors |

The ZIKV-SWITCH and WNV-SWITCH reporters were introduced into A549 cells, which were then infected with ZIKV or WNV, respectively (MOI 5). Similar to our validation strategy with DENV-SWITCH, we compared the activation of the reporters with the detection by immunofluorescence of the NS2B and E proteins for ZIKV and WNV, respectively. The percentages of infected cells detected by the reporters and by NS2B/E immunostaining did not differ significantly in the matched comparisons (Fig. 4B). To determine the potential of these reporters to evaluate antiviral compound activity, we performed a compound assay with MK0608 and Niclosamide on ZIKV and WNV infections. Both compounds reduced reporter-defined infection in a dose-dependent manner. Absolute IC50 values for MK0608 were 7.22 µM for ZIKV and 6.55 µM for WNV. Those for niclosamide were 3.21 µM for ZIKV and 1.44 µM for WNV (Fig. 4C–D). To further evaluate ZIKV-SWITCH, we determined infection efficiency in the context of EMC6 and RACK1 silencing, both previously described as ZIKV host factors required for infection ^24,46^. RACK1 silencing reduced ZIKV-SWITCH activation by 84.2% (P = 0.015). EMC6 silencing produced a 32.4% higher mean response, but this difference was not statistically significant (P = 0.2886; Fig. 4E). To confirm this observation with a reporter-independent readout and mitigate the cell background differences, we transfected the same siRNA into HeLa cells and quantified intracellular ZIKV RNA by RT-qPCR. ZIKV RNA abundance was also higher following EMC6 silencing in two independent experiments (Sup Fig. 7).

To validate V-SWITCH in a different viral family, we substituted the PCS with a sequence recognized by the coronavirus OC43 protease nsp5/3CL. We introduced OC43-SWITCH into a HEK293T cell line and infected it with OC43 (MOI 0.25) for 48 hours before detecting the infection by immunostaining and flow cytometry. At 48 hpi, the mean percentages of positive cells were 77.9% by anti-OC43 immunostaining and 59.1% by OC43-SWITCH fluorescence. Reporter fluorescence detected a lower fraction of cells in the infected condition and showed higher background in the uninfected condition than immunostaining (P = 0.0264 for each comparison; Fig. 4F). To evaluate the utility of the reporter for antiviral testing, we treated OC43-infected cells with Ferrostatin, a ferroptosis inhibitor, recently reported to have antiviral activity against OC43 ^47^. Ferrostatin treatment (60 µM) reduced OC43-SWITCH activation by 54.5% reduction (P = 0.0159; Fig. 4G). These results establish the V-SWITCH reporter architecture as a flexible platform that can be rapidly reprogrammed to detect the activity of diverse viral proteases and deployed across multiple cell lines.

## Discussion

Real-time, single-cell monitoring of viral infection remains a critical need for both mechanistic virology and screening and characterization of antiviral targets. Here, we describe V-SWITCH, a single-vector, host-encoded reporter that converts viral protease activity during infection into an OFF-to-ON fluorescent signal. The release-and-capture architecture of this reporter combines an ER-anchored mNG3A(1–10) fragment with a nuclear mNG(11)-BFP capture module, providing low background, high specificity, and compatibility with live-cell imaging and flow cytometry-based methods.

We developed V-SWITCH to complement existing reporters and open new experimental options. Our reporter detects protease activity during live-virus infection, with a high signal to noise ratio, and enables quantification and selection of infected populations. These capabilities build on earlier protease reporters, including those using alternative strategies for activation, i.e fluorescence quenching, validated using ectopic protease expression or non-replicating surrogates, and translocation-based reporters which redistribute an existing fluorescent signal and are highly suited to microscopy approaches ^10–14^.

V-SWITCH was also designed for modularity and adaptability. Because both reporter modules are encoded in a single lentiviral vector, the system does not require endogenous cell or virus tagging and facilitates deployment across cell types and viruses, as shown in A549, BJ-5ta, and HEK293T cells and with DENV, ZIKV, WNV, and HCoV-OC43. Also, the host cell encodes V-SWITCH, so it can be adapted to a broad range of protease-expressing RNA viruses without modifying their genomes or compromising fitness, overcoming a key limitation of systems depending on reverse genetics. This feature is especially valuable for studying unmodified, low-passage clinical isolates, which remain difficult to investigate using conventional reporter viruses ^10,48,49^. V-SWITCH could therefore facilitate comparative studies across viral families, strains, and clinical isolates, provided that the selected cells support viral replication and a suitable protease-cleavage sequence is available.

V-SWITCH makes it possible to track and study viral infection dynamics at the single cell level. A constitutive BFP signal reports reporter expression, and an mNG signal reports protease activity. Baseline BFP fluorescence was strongly correlated to mNG fluorescence, underscoring the need to consider reporter expression when interpreting signal strength. Live-cell imaging with DENV-SWITCH revealed substantial cell-to-cell heterogeneity in reporter activation and fluorescence intensity. Among cells whose response timing could be reliably estimated, low-, medium- and high-reporter activation responses reached half of their fluorescence increase at similar median times within each replicate. In an independent sorting experiment, populations with higher mNG/BFP ratios contained more intracellular DENV RNA. These findings support the use of V-SWITCH not only to identify infected cells, but also to distinguish populations with different levels of viral RNA accumulation. Heterogeneous infection outcomes have been associated with differences in cell state, innate immune activity, cell-cycle stage, metabolism, and stochastic variation in viral uptake and replication ^50,51^. By combining longitudinal imaging with the isolation of defined cell populations, V-SWITCH provides a means of investigating why some cells accumulate more viral RNA than others. Coupling reporter-based selection with transcriptomic, proteomic, or optical pooled-screening approaches could help identify the cellular states and host factors that favor or restrict infection ^52,53^.

The correspondence between fluorescence intensity and viral RNA enabled V-SWITCH to support pooled CRISPRi screening. A CRISPRi screen using a druggable sgRNA library identified 29 statistically significant genes, and we prioritized nine high-confidence candidates for experimental follow-up. The screen recovered factors previously associated with DENV or flavivirus infection, including DDOST, ATP6AP1, SRPR, and EEF1A1, while identifying TAF1, GTF2A1, DNMT1, and FEN1 as less-characterized DENV dependencies in human cells ^40–42^. Together, the prioritized candidates implicate ER protein targeting and processing, organelle acidification, translation, transcriptional and epigenetic regulation, phosphoinositide metabolism, and DNA maintenance in efficient DENV infection. Validation by siRNA-mediated depletion reduced infection by 75–99% providing independent functional validations of their pro-viral role.

We used V-SWITCH to measure the anti-DENV activity of inhibitors against three targets: the ER chaperone BiP/GRP78, which we selected from the literature, and the translation elongation factor EEF1A and the phosphatidylinositol 4-kinase PI4KA, which we selected from our CRISPRi screen. HA15 is a BiP/GRP78 ATPase inhibitor with known activity against ZIKV and several dsDNA viruses ^26–28^. In BJ-5ta fibroblasts, HA15 reduced DENV infection without a detectable reduction in cell viability, an observation consistent with the upregulation of GRP78 during DENV infection. The EEF1A inhibitor Plitidepsin was active against DENV at a sub-nanomolar IC50, which to our knowledge, extends its reported antiviral spectrum to DENV and supports the EEF1A dependency identified in our screen. The PI4KA inhibitor GSK-A1 had a weaker effect on DENV infection than PI4KA silencing. This difference may reflect the mode of perturbation, since GSK-A1 inhibits catalytic activity whereas siRNA depletion removes the protein and its non-catalytic functions, or the difference in cell type (A549 for silencing, BJ-5ta for GSK-A1) ^54^. The activity of GSK-A1 is notable given earlier reports of little PI4KA requirement in DENV reporter-virus and replicon systems, and it raises the possibility of a context-dependent role for PI4KA ^44^. Together, these results extend the antiviral activity of HA15 and Plitidepsin to DENV and show that V-SWITCH can support the identification of antiviral targets.

The application of V-SWITCH to multiple flaviviruses also allowed us to compare their dependence on shared host factors. Whereas EMC6 silencing reduced DENV reporter activation, it was associated with higher ZIKV reporter activation in A549 cells, although this increase was not statistically significant. Higher ZIKV RNA abundance was also observed in two independent experiments in parental HeLa cells. These preliminary observations differ from previous studies identifying EMC6 as a ZIKV dependency factor ^46^. Differences in the strength of EMC6 depletion, infection MOI and duration, and cell background could contribute to these different outcomes. In particular, acute silencing and prolonged depletion may affect the wider EMC differently ^55^. Beyond supporting viral-protein biogenesis, the EMC has also been linked to ER–mitochondria communication ^56^. Freppel and colleagues showed that disrupting these contacts through depletion of other tethering proteins increased viral replication and reduced ZIKV-induced apoptosis ^57^. Altered ER–mitochondria communication therefore provides another possible explanation for the increased ZIKV readouts following EMC6 silencing. However, whether EMC6 depletion affects these contacts in our experimental conditions remains unknown, and the apparent increase in ZIKV infection requires further validation.

Some limitations of V-SWITCH should be considered. This reporter detects viral protease activity and therefore does not measure viral entry, genome replication, particle assembly, or production of infectious progeny. Activation requires sufficient accumulation of active protease, producing a delay relative to the early stages of infection. Split-fluorescent complementation is also irreversible, resulting in cumulative fluorescence rather than an instantaneous measurement of protease activity. Although gating on the constitutive BFP module controls for heterogeneous reporter expression, perturbations affecting transcription, translation, protein folding, or protein stability could influence reporter abundance independently of infection. Hits from reporter-based screens should therefore be confirmed using reporter-independent measurements such as viral RNA, viral-protein staining, or infectious-virus production. Finally, the CRISPRi screen was restricted to a druggable-gene library and a single A549 cell background; viral dependencies may differ among cell types, viral strains, and clinical isolates.

In summary, V-SWITCH combines the sensitivity of split-fluorescent complementation with a modular, host-encoded architecture that supports live-cell imaging, quantitative flow cytometry, cell sorting, antiviral testing, and pooled functional-genomic screening. Its adaptability to diverse protease-expressing RNA viruses without requiring viral-genome modification provides a flexible framework for studying replication heterogeneity and discovering host dependencies. This advantage may be particularly valuable for genetically diverse or low-passage clinical isolates for which reporter viruses and reverse-genetic tools remain limited.

## Methods

### Cell culture

A549 (ATCC, CCL-185), HEK 293T (ATCC, CRL-3216), HEK 293FT (Invitrogen, R70007), and BJ-5ta (ATCC, CRL-4001) cell lines were maintained in Dulbecco’s modified Eagle’s medium (DMEM; Gibco) supplemented with 10% foetal bovine serum (FBS; Gibco) at 37°C in a humidified atmosphere containing 5% CO₂. Cells were passaged every 2–3 days upon reaching approximately 80% confluence and routinely tested for mycoplasma contamination. All cell lines were used within 20 passages of thawing.

### Virus production and titration

DENV-2 stocks (ATCC, VR-1584) were propagated using a Vero–C6/36 co-culture system under antibiotic-free conditions. Vero cells were adapted to 28°C by passaging at least twice at reduced temperature before co-culture setup. Vero and C6/36 cells were combined at a 1:4 ratio and seeded in T175 flasks containing DMEM supplemented with 10% FBS. Co-cultures were incubated at 28°C with 5% CO₂ until reaching 90% confluency.

Infections were performed at low multiplicity of infection (MOI ∼0.001–0.01) using virus stocks previously titrated on A549 cells. Infected co-cultures were incubated at 28°C and supernatants harvested when cytopathic effects were evident (typically 5–7 days post-infection). Viral supernatants were clarified by centrifugation at 300×g for 10 minutes, filtered through 0.45 μm filters, aliquoted, and stored at −80°C.

Viral titers were determined by plaque assay on A549 cells. Cells were seeded in 24-well plates and infected with serial dilutions of virus stock. After 1-hour adsorption, cells were overlaid with 1% methylcellulose in DMEM and incubated at 37°C for 48–72 hours. Cells were fixed with 4% paraformaldehyde, permeabilized with Triton X-100, blocked with 1% BSA, and stained with anti-flavivirus E protein antibody (4G2) followed by Alexa Fluor 488-conjugated secondary antibody. Plaques were counted by fluorescence microscopy and titers expressed as plaque-forming units (PFU) per milliliter.

Zika virus (ZIKV-PRVABC-59 strain-NCBI: KX377337), and West Nile virus (WNV-NY99 strain - NCBI: AY842931) stocks were propagated as previously described ^59,60^. Briefly, Vero or C6/36 cells were infected at a low multiplicity of infection (MOI 0.01–0.1) and viral supernatants were collected at peak cytopathic effect, clarified by centrifugation (3,000 × g, 10 min, 4°C), filtered through a 0.45 µm membrane, aliquoted, and stored at −80°C. Viral titres were determined by plaque-forming assay on Vero cells.

Human coronavirus OC43 (HCoV-OC43) was produced and titrated as described by Hein et al. (2025).

### Viral infection assays

For DENV, ZIKV, and WNV infection assays, 1 × 10⁵ cells were seeded per well in 24-well plates in 500 µL of complete medium and allowed to adhere overnight. The following day, cells were infected at the indicated MOI (typically MOI 5 unless otherwise stated). Viral inoculum was added directly to the well and removed after 1 hour. Cells were washed once with DMEM and maintained in fresh DMEM supplemented with 10% FBS for the indicated time periods (typically 24 h) before analysis. An uninfected control was included in all experiments. For HCoV-OC43 infections, HEK 293T cells were infected at MOI 0.25 and maintained for 48 h before fixation and analysis.

### Plasmid construction

Cleavable Anchor Module (CAM) and Nuclear Capture Module (NCM) sequences were designed as synthetic gene blocks (gBlocks; Twist Bioscience). For the first-generation reporter, the CAM comprised, from N- to C-terminus: a nuclear localisation signal (NLS), the mNG3A(1– 10) fragment, a viral protease cleavage site (PCS), the transmembrane domain of Sec61β (ER anchor), a T2A self-cleaving peptide, and a blasticidin resistance gene, all driven by the EF1α promoter. The NCM was generated by CRISPR-based knock-in of the mNG(11) fragment at the endogenous NPM1 or HNRNPC locus using a single-stranded oligodeoxynucleotide (ssODN) donor and Cas9/gRNA ribonucleoprotein delivery as described in Cho et al., 2022.

For the second-generation single-vector reporter (V-SWITCH), both modules were encoded on a single lentiviral backbone. The CAM (EF1α promoter) retained the NLS–mNG3A(1–10)–PCS– ER anchor–T2A–blasticidin architecture, while the NCM (CMV promoter) expressed an NLS– mNG(11)–ALFA-tag–TagBFP–NLS fusion. The NCM was designed based on a mNG(11)-BFP sequence provided by Bo Huang (UCSF) and modified to express the dimmer TagBFP. The mNG3A(1–10) variant was used to enhance complementation efficiency. The BFP fusion served as a constitutive reporter expression marker and nuclear segmentation channel.

For adaptation to additional viruses, PCS cassettes encoding recognition sequences for the ZIKV NS2B/3 protease, WNV NS2B/3 protease, or HCoV-OC43 nsp5/3CL protease were synthesised as complementary oligonucleotides with NotI and BamHI overhangs. Oligos were annealed by mixing 1 µL of each strand (100 µM) with 25 µL of 2× annealing buffer (200mM Potassium Acetate, 60mM HEPES, 4mM Magnesium Acetate) and 23 µL water, heating to 95°C for 5 min, and cooling at −1.2°C min⁻¹ for 60 cycles. Annealed duplexes were diluted 1:200 prior to ligation.

HCoV-OC43 nsp5/3CL protease cleavage site was obtained by accessing the genome of HCoV-OC43 strain 1588 through the *Ensembl gene browser* and obtaining an amino acid sequence flanking nsp14/nsp15 previously characterized to be efficiently cleaved by nsp5/3CLpro in a SARS-CoV-2 reporter system ^12^.

Restriction digestions were performed in 10 µL reactions containing 1 µg plasmid DNA or 250 ng gBlock, 0.5 µL of each restriction enzyme, and 2 µL of 10× CutSmart buffer (NEB), with nuclease-free water to volume. Reactions were incubated for 1 h at 37 °C. Digested backbones were resolved on 1% agarose gels and extracted using the Zymoclean Gel DNA Recovery Kit (Zymo Research). Inserts were purified using the DNA Clean and Concentrator Kit (Zymo Research) and eluted in 6 µL elution buffer.

Ligation reactions were assembled in 10 µL total volume containing 50 ng backbone DNA, insert at 5:1 ratio (calculated using the NEBioCalculator), 1 µL T4 DNA Ligase Buffer, and 1 µL T4 DNA Ligase (NEB), and incubated at 25°C for 1 h. Ligation products (2.5 µL) were transformed into 50 µL of STBL3 chemically competent cells (Thermo Fisher) by incubation on ice for 10 min, heat shock at 42°C for 45 s, and recovery in 200 µL SOC medium at 37°C for 1 h with shaking at 300 rpm. Transformations were plated on LB agar containing carbenicillin and incubated overnight at 37°C. Single colonies were inoculated into 2.5 mL LB with carbenicillin and grown overnight, and plasmid DNA was extracted using the ZymoPURE Plasmid Miniprep Kit (Zymo Research). All constructs were verified by Sanger sequencing.

The DENV-SWITCH, ZIKV-SWITCH, WNV-SWITCH, and OC43-SWITCH plasmids have been deposited with Addgene under accession numbers 262011, 262012, 262013, and 262014, respectively.

### Lentivirus production

Lentiviral particles were produced using a third-generation packaging system in HEK 293FT cells. On Day 1, 2 × 10⁶ HEK 293FT cells were seeded per T25 flask in DMEM GlutaMAX supplemented with 10% FBS, aiming for 60–80% confluence the following day. On Day 2, transfection complexes were prepared as follows. For each flask, the packaging DNA mix contained 370 ng pMD2.G (VSV-G envelope), 370 ng pRSV-Rev, 740 ng pMDLg/pRRE, 240 ng pAdVAntage, and 570 ng transfer plasmid, brought to 10 µL with nuclease-free water. In parallel, 13 µL of Lipofectamine 2000 (Invitrogen) was diluted in 750 µL of room-temperature Opti-MEM I (Invitrogen), mixed gently, and incubated for 5 min. The packaging DNA mix was diluted separately in 750 µL of room-temperature Opti-MEM I. The DNA–Opti-MEM solution was then added dropwise to the Lipofectamine–Opti-MEM solution, mixed gently, and incubated for 20 min at room temperature. One millilitre of warm Opti-MEM I was added to the transfection complex, the culture medium was aspirated from the 293FT cells, and the resulting 2.5 mL of transfection mix was carefully dispensed along the side of the flask to minimise cell detachment. At 5–6 h post-transfection, the transfection medium was replaced with 5 mL of fresh DMEM supplemented with 10% FBS. At 24 h post-transfection, the medium was replaced again with 5 mL of DMEM supplemented with 10% FBS. At 48 h post-transfection, viral supernatants were collected, clarified by centrifugation (500 g, 5 min, room temperature), and filtered through a 0.45 µm low-protein-binding filter. Lentiviral stocks were used immediately for transduction or aliquoted and stored at −80°C.

### Generation of stable reporter cell lines

A549, BJ-5ta, or HEK 293T cells were seeded in 6-well plates at approximately 60% confluence. The following day, cells were transduced with 1 mL of lentiviral supernatant per well and incubated at 37°C for 24 h. Viral supernatant was then removed, cells were washed twice with PBS, and maintained in complete medium for an additional 48 h before initiating selection with 5 µg/mL blasticidin. Surviving cells were expanded and subjected to fluorescence-activated cell sorting (FACS) to isolate single-cell clones with homogeneous BFP reporter expression levels.

### Flow cytometry and immunofluorescence

Infected and control cells were harvested by trypsinisation, transferred to round-bottom 96-well plates, and fixed with 4% paraformaldehyde (PFA) at room temperature for 15 min. For immunofluorescence staining, fixed cells were centrifuged at 500 × g for 5 min, permeabilized in BD Perm/Wash Buffer (BD Biosciences, 554723) for 30 min, and incubated with primary antibody (1:1,000 dilution in Perm/Wash Buffer) for 1 h at room temperature. After two washes with PBS, cells were incubated with Alexa Fluor 555-conjugated secondary antibody (1:1,000 dilution - anti-rabbit-AF555 Invitrogen, A32732 or anti-mouse-AF555 Invitrogen, A32727) for 1 h, then washed twice with Perm/Wash Buffer, and resuspended in PBS. The following primary antibodies were used: anti-DENV NS3 (Genetex, GTX124252), anti-ZIKV NS2B (Genetex, GTX133308), anti-WNV E protein (Novus, Nbp2-52709), and anti-HCoV-OC43 nucleoprotein (Millipore Sigma, mab9012). Samples were analysed on a CytoFLEX flow cytometer (Beckman Coulter). Flow cytometry data were processed using FlowJo software (v10; BD Biosciences).

### Antiviral compound assays

For DENV, ZIKV, and WNV compound assays, reporter cell lines were seeded in triplicate per condition the day before the experiment. Cells were pre-treated with serial dilutions of MK0608 (Sigma Aldrich, SML3200), niclosamide (MedChemExpress, HY-B0497), GSK-A1 (MedChemExpress, HY-125118), or Plitidepsin (MedChemExpress, HY-16050) for 2 h, alongside a dimethyl sulfoxide (DMSO - Sigma Aldrich, D26500) vehicle control. Cells were then infected at MOI 5 in the absence of compound for 1 h. After removal of the viral inoculum, compound-containing medium was restored. Cells were fixed with 4% PFA at 24 h post-infection and analysed by flow cytometry.

For HCoV-OC43 compound assays, HEK 293T OC43-SWITCH cells were infected at MOI 0.25 in the presence of ferrostatin-1 (60 µM) or DMSO vehicle control. Cells were fixed at 48 h post-infection and analysed by flow cytometry.

For the evaluation of the BiP/GRP78 inhibitor HA15, BJ-5ta DENV-SWITCH cells were treated with HA15 at 50 µM for 2 h before being infected with DENV (MOI 5) for 1 h. The viral inoculum was then removed, and HA15 or DMSO-containing medium was restored for 24 h before fixation and flow cytometric analysis.

Inhibition was quantified by normalising the percentage of reporter-positive cells in each condition to the DMSO-treated control (set to 100%). Absolute half-maximal inhibitory concentrations (IC50) were defined as the fitted compound concentrations giving 50% of the DMSO-control response. Normalised infection-response data were fitted using the logistic inhibition model described under Statistical analysis.

Cell viability was measured by measurement of LDH release using CytoTox-ONE™ Homogenous Membrane Integrity Assay (Promega, G7890) with values normalized using DMSO and plate-matched cell-lysate controls, set to 100% and 0%, respectively. Fluorescence was read using a SpectraMax plate reader (Molecular Devices).

### siRNA-mediated gene silencing

A549 V-SWITCH cells were seeded at 3,000 cells per well in 96-well plates in 100 µL of complete growth medium and allowed to adhere overnight. The following day, siRNA transfections were performed in triplicate. For each well, 0.25 µL of Lipofectamine RNAiMAX (Thermo Fisher) was combined with 0.25 µL of 10 µM siRNA in 25 µL Opti-MEM, and the mixture was incubated at room temperature for 10 min. The transfection complex (25 µL) was then added directly to each well. At 72 h post-transfection, cells were infected with DENV or ZIKV at MOI 5 for 24 h, fixed with 4% PFA, and analysed by flow cytometry. The siRNA references can be found in Supplementary Table 1. Gene silencing efficiency was confirmed by RT-qPCR or western blot (Sup. Fig. 4 and 6). RT-qPCR primer sequences are provided in Supplementary Table 2.

### Live-cell imaging and time-lapse microscopy

A549 DENV-SWITCH cells were seeded at 50,000 cells per well in glass-bottom 24-well plates and allowed to adhere overnight. Cells were infected with DENV or ZIKV at MOI 5 for 1 h, after which the inoculum was removed and replaced with FluoroBrite DMEM (Gibco) supplemented with 10% FBS. Time-lapse imaging was performed on a confocal microscope at one-hour intervals. The two biological experiments were analyzed over their 27 frames (1.5–27.5 hpi). Phase images were computationally reconstructed using waveorder (https://github.com/mehta-lab/waveorder).

### Image analysis pipeline

#### Cell tracking and fluorescence measurements

Nuclei were segmented in the constitutive BFP channel using Cellpose (v4.0.7; Stringer et al., 2021) and tracked using Ultrack (Bragantini et al., 2025). The same analysis was applied to two independent biological experiments, using 27 consecutive frames from each (frames 0–26; 1.5–27.5 hpi). Tracks with at least six consecutive observations were retained. Mean nuclear mNG and BFP intensities were measured at each time point, and their ratio was calculated.

#### Reporter activation

For each replicate, the activation threshold was the mean mNG/BFP ratio plus three standard deviations across all measurements in the matched uninfected control well. Reporter activation required two consecutive frames above this threshold; the first defined the detection time. Tracks already meeting this criterion at their first recorded frame were excluded because the transition to reporter positivity had not been observed. Because cells were tracked for different portions of the movie, the cumulative proportion becoming reporter-positive was estimated using Kaplan–Meier analysis. Each cell entered at its first tracked frame and was followed until activation.

#### Baseline and maximum fluorescence

These analyses included 724 cells with valid mNG and BFP measurements at all six baseline frames (frames 0–5; 1.5–6.5 hpi) and a detected reporter activation after this baseline period. Baseline fluorescence was averaged across these frames; maximum mNG fluorescence was the highest measured value during each cell’s track. The same cells were used in both panels. Each intensity measure was divided by its median among eligible cells in the corresponding replicate, then log2-transformed before combining replicates.

#### Response magnitude and timing

We fitted a curve to each cell’s mNG/BFP measurements to estimate its final response level and when it reached half of this response. Among the 2,923 eligible cells, the lowest, middle and highest thirds of estimated response levels within each replicate were classified as Low, Medium and High activators.

We compared timing only in cells whose measurements supported a reliable estimate of t50 with a fitted curve R² ≥ 0.7 and timepoints covering the entire activation. This retained 626 cells (21.4% of the initial group). Figures 2E–F show these same cells, keeping their original Low, Medium and High assignments. Response levels were expressed relative to the median of all initially eligible cells in each replicate, and timing was shown separately by replicate.

### FACS sorting and RT-qPCR

A549 DENV-SWITCH cells were seeded at 500,000 cells per well in 6-well plates and allowed to adhere overnight. Cells were infected with DENV at MOI 5 for 1 h, the inoculum was removed, and cells were maintained in complete medium for 24 h. Cells were then detached by trypsinisation, pooled (two wells per replicate), and passed through a cell strainer to remove clumps. Cells were sorted on a Bigfoot cell sorter (Thermo Fisher) into four populations based on the mNG/BFP fluorescence intensity ratio: mNG-negative, low mNG, mid mNG, and high mNG. For each gate, 10,000 cells were collected directly into 100 µL of RNA Lysis Buffer (Zymo Research).

Total RNA was extracted and reverse-transcribed, and DENV viral RNA levels were quantified by RT-qPCR using virus-specific primers (Supplementary Table 2). Viral RNA abundance was normalised to GAPDH mRNA as an internal reference. For plotting, technical-replicate 2^−ΔCq values were averaged within each biological sample, divided by the arithmetic mean of the mid-mNG biological-sample values, and multiplied by 100. Statistical inference used paired ΔCq values calculated after averaging technical Cq measurements.

### Design of the druggable-target CRISPRi library

Genes encoding targets classified as Tchem or Tclin in the Pharos database were selected to generate a druggable-target CRISPRi library ^62^. sgRNA sequences targeting these genes were drawn from the Top5 subset of the human CRISPRi-v2 library ^63^. The final library comprised 14,545 sgRNAs, including 14,214 gene-targeting sgRNAs covering 2,583 genes and 331 non-targeting control sgRNAs.

### sgRNA library cloning and lentiviral production

The pooled oligonucleotide library was synthesized by Twist Bioscience and amplified by dial-out PCR ^64^ using KAPA HiFi HotStart ReadyMix and the primers listed in Supplementary Table 2. The amplified oligonucleotides were cloned into the PerturbView lentiviral vector ^65^ by Golden Gate assembly using Esp3I and T4 DNA ligase. The assembled library was transformed into MegaX electrocompetent cells and packaged into lentiviral particles using the LV-MAX Lentiviral Production System (ThermoFisher). Library representation was maintained at greater than 1,000-fold coverage throughout cloning, bacterial expansion, and lentiviral production. The sgRNA sequences and their gene target are listed in Supplementary Table 3.

### Pooled CRISPRi screening and cell sorting

A549 DENV-SWITCH cells were transduced with a lentiviral vector encoding ZIM3-KRAB-dCas9–mCherry (Addgene #154473). Cells displaying the highest 10% of mCherry fluorescence were isolated by FACS and expanded to generate the screening population. Cells were subsequently transduced with the pooled druggable-target CRISPRi sgRNA lentiviral library at MOI 0.2. Following transduction and selection, cells were expanded while maintaining at least 1000-fold library representation.

Cells were infected with DENV at MOI 5 and sorted at 24 h post-infection. Within the population strongly expressing the constitutive BFP reporter module, cells were separated into a Low population comprising the 10% of cells with the lowest V-SWITCH activation and a Top population comprising the 10% with the highest activation. The screen was performed in two independent biological replicates. Infected unsorted cells were collected in parallel for quality control but were not included in the Top-versus-Low analysis.

### Preparation of sgRNA libraries for sequencing

Genomic DNA was extracted from the screened cell populations using the Quick-DNA Miniprep Plus Kit (Zymo Research, D4069). For each sample, sgRNA sequences were amplified from 40 µg of genomic DNA using a two-step PCR with KAPA HiFi HotStart ReadyMix. PCR1 was performed for 18 cycles. Indexed Illumina adapters were added by PCR2 for 12 cycles. Cycling conditions were 95°C for 3 min, followed by the indicated number of cycles of 98°C for 20 s, 67°C for 20 s and 72°C for 30 s, with a final extension at 72°C for 5 min. Amplified libraries were pooled, purified at a 1:1 ratio using SPRIselect beads (Beckman Coulter), and quantified using a Qubit fluorometer. Libraries were sequenced on an Illumina NextSeq 2000. Sequencing generated 51–128 million reads per sample, of which 96.3% mapped to the sgRNA reference. PCR1, PCR2, and custom sequencing primers sequences are available in Supplementary Table 2.

### CRISPRi screen analysis and hit selection

Sequencing reads were assigned to sgRNA spacers using MAGeCK count v0.5.9.5 with a spacer length of 19 nucleotides and the Druggable Targets library reference ^66^. The resulting raw count matrix contained 14,545 sgRNAs across six samples: Unsorted, Top, and Low populations from each of two biological replicates. Of these guides, 14,214 targeted 2,583 genes and 331 were NTCs.

The Top-versus-Low comparison was performed using the maximum-likelihood module of MAGeCK v0.5.9.5 ^67^. Only the four sorted samples (Low_1, Low_2, Top_1, and Top_2) were included. The design matrix contained a common baseline term and a Top-bin indicator, with the Low samples serving as the reference. The calculated Z-score therefore represented enrichment or depletion in the Top population relative to the Low population. Negative Z-scores indicated that downregulation of the corresponding gene shifted cells towards the Low reporter population.

Counts were normalized using the MAGeCK median-normalization method. MAGeCK-MLE modeled sgRNA counts using a negative binomial distribution and estimated gene-level beta coefficients by expectation maximization. The option --max-sgrnapergene-permutation 11 was used, excluding SGK3, ESRRG, and the aggregated negative-control entry and leaving 2,581 targeted genes with estimated Z-scores.

Statistical significance was assessed using MAGeCK’s guide-count-matched permutation procedure with 100 permutation rounds. Permutation-derived P values were adjusted for multiple testing using the Benjamini–Hochberg procedure, and the permutation FDR reported in the Top|fdr output column was used for hit calling. Genes with permutation FDR < 0.05 were classified as screen hits. For experimental follow-up, hits were additionally required to have |Z| > 5, where Z is the Wald statistic calculated by MAGeCK-MLE as the beta coefficient divided by its standard error. This prioritization yielded nine candidates, all with negative Z-scores. The CRISPRi screen analysis results are presented in Supplementary Table 4.

### mNG(1–10) and mNG11 protein stability assay

To assess the relative stability of the split mNG fragments, cells were transiently transfected with plasmids encoding either mNG(1–10) (Addgene #82610), 7 tandem repeats of mNG11 fused to a BFP (a gift from Bo Huang), or a full length mNG (a gift from Manuel Leonetti). A549 cells were seeded in a 24 well plate at 50,000 per well and left to adhere overnight. Then, the cells were transfected with 500ng of plasmid DNA and 1 µL of Lipofectamine 2000 (Invitrogen), following the manufacturer protocol. Percent fluorescent cells were analyzed by flow cytometry at 72 hours post-transfection.

### Statistical analysis

Statistical analyses were performed using GraphPad Prism 11; revised imaging cohort selection, survival summaries and reporter-timing validation were performed using Python. Unless otherwise stated, data are presented as mean ± standard deviation (s.d.) from three independent biological experiments. Technical replicates were averaged to generate one value per biological experiment. Measurements obtained within the same independent experiment were treated as matched observations. All statistical tests were two-sided, with α = 0.05. No replicate-level outliers were excluded.

For comparisons involving two matched conditions, paired two-tailed t-tests were used. When multiple paired comparisons constituted one prespecified family, P values were adjusted using the Holm–Šídák method. Comparisons of three or more matched conditions were performed using repeated-measures one-way ANOVA followed by Dunnett’s multiple-comparisons test against the indicated control. Grouped experiments containing two matched factors were analyzed using two-way repeated-measures ANOVA followed by Šídák-adjusted comparisons between detection methods within each infection condition. For effects involving factors with more than two levels, sphericity was not assumed and Geisser–Greenhouse corrections were applied. Because most experiments contained three biological replicates, normality was not formally tested. Parametric analyses were performed using replicate-level continuous measurements, ΔCq values for RT–qPCR experiments, or log10-transformed fluorescence-intensity measurements where indicated.

For RT–qPCR, technical replicates were averaged before ΔCq calculation relative to GAPDH. Inference was performed on paired ΔCq values. For plotting, relative expression was calculated as 2^−ΔCq, divided by the arithmetic mean of the corresponding control 2^−ΔCq values, and multiplied by 100. In Fig. 2G, technical-replicate 2^−ΔCq values were averaged before normalization to the mean mid-mNG control; inference still used paired ΔCq values calculated from averaged technical Cq measurements.

Infection concentration–response data were fitted by unweighted ordinary least-squares nonlinear regression using Prism’s “Absolute IC50, X is concentration” variable-slope logistic inhibition model. Technical replicates were averaged within each independent biological experiment, yielding one value per biological experiment at each concentration; a single curve was fitted to all biological-replicate values. For DMSO-normalized responses, Top was fixed at 100%, the reference Baseline was fixed at 0%, Bottom was fitted subject to a non-negative constraint, and absolute IC50 was positive. Baseline defines the 50% reference and is distinct from the fitted Bottom plateau. Thus, absolute IC50 denotes the concentration giving 50% of the DMSO-control response. Zero-concentration DMSO controls were included in the fits but omitted from logarithmic concentration axes. Profile-likelihood 95% confidence intervals were reported when estimable. The niclosamide estimate in Fig. 1G is approximate, with no estimable 95% interval; the upper confidence limit for niclosamide in the HNRNPC-NCM line in Supplementary Fig. 1H could not be determined. No absolute IC50 was reported for GSK-A1 because the mean response did not reach 50% of DMSO within the tested range. Cell-viability measurements in concentration–response panels were displayed as measured points with error bars without fitted inhibition curves; no CC50 was estimated where viability did not approach 50%. In Fig. 4G, both control and treated replicate values were divided by the arithmetic mean of the three DMSO biological controls and multiplied by 100; inference used the three matched biological-experiment pairs.

Single-cell imaging data were obtained from two independent biological experiments and were analyzed descriptively. Associations were summarized using Spearman’s rank correlation coefficient (ρ). Correlations for combined, within-replicate-normalized Figure 2C–D data are cell-weighted descriptive summaries, not replicate-adjusted inference. Timing associations were summarized separately by biological replicate. No inferential P values or biological-replicate confidence intervals are reported for these cell-level correlations or imaging-derived group comparisons. Curve-fit profile intervals used for timing-quality selection quantify conditional fit uncertainty only.

Supplementary Fig. 2B shows both independent biological experiments and was treated descriptively. Supplementary Figs. 3 and 5 are representative gating illustrations and were not statistically analyzed. Supplementary Fig. 7 was based on two paired independent biological experiments and was summarized descriptively without inferential testing. Exact sample sizes are provided in the corresponding figure legends. Exact P values, test statistics, degrees of freedom, effect estimates and 95% confidence intervals are provided in the Source Data. For figure annotations, *P < 0.05, **P < 0.01, ***P < 0.001 and ****P < 0.0001.

## Data availability

The raw and processed sequencing data generated in the pooled CRISPRi screen have been deposited in the NCBI Gene Expression Omnibus (GEO) under accession number GSE343702. All other data supporting the findings of this study are available within the article and Supplementary Information or from the corresponding authors upon reasonable request. Source data are provided with this paper.

## Code availability

The image-analysis code used for Figure 2 (version 2.0.0) and the MAGeCK-based CRISPRi screen analysis code used for Figure 3 (version 1.0.0) are archived on Zenodo under DOIs 10.5281/zenodo.23194567 and 10.5281/zenodo.23194649, respectively. Access to both deposits is currently restricted pending review for public release; requests for access should be directed to the corresponding authors. The associated GitHub repositories, VSWITCH-reporter-analysis and denv-crispri-screen, are private. Code development was assisted by Claude Opus (Anthropic) and Codex (OpenAI).

## Contributions

M.C.-R., V.T.-L., and C.A. conceptualized the project. M.C.-R., V.T.-L., and M.G. developed the methodology. M.C.-R., M.G., A.M., S.-C.L., H.W., S.P., S. Kanwar, S. Khadka, and D.J. performed experiments. V.T.-L. developed the image analysis software with code review from S.P. V.T.-L. processed and analyzed the CRISPRi screen NGS data. M.C.-R., V.T.-L., and S.P. analyzed data. S. Khadka and A.M. provided key reagents. M.C.-R. and V.T.-L. prepared figures and wrote the manuscript. C.A. wrote and reviewed the manuscript and figures. S.M., R.H., V.T.-L., and C.A. supervised the work.

## Acknowledgements

We thank Manuel Leonetti, Rodrigo Baltazar-Nuñez, Camille Januel, and Madhurya Sekhar for providing the HCoV-OC43 translocation reporter plasmid backbone and the reagents used for CRISPR knock-in of mNG(11) in NPM1 and HNRNPC, Bo Huang for providing the plasmid encoding mNG(11)-BFP used to design the V-SWITCH NCM module, Aofei Liu for code review of the image analysis pipeline, and Mosopefoluwa John for experimental support during his internship. Schematics in Figures 1, 2, and 4 were created in BioRender. We thank the Biohub and its donors Priscilla Chan and Mark Zuckerberg for funding this work.

