## Supplementary_Figures_1-7 for "V-SWITCH: A single-vector OFF-to-ON fluorescent reporter of live RNA virus infections"

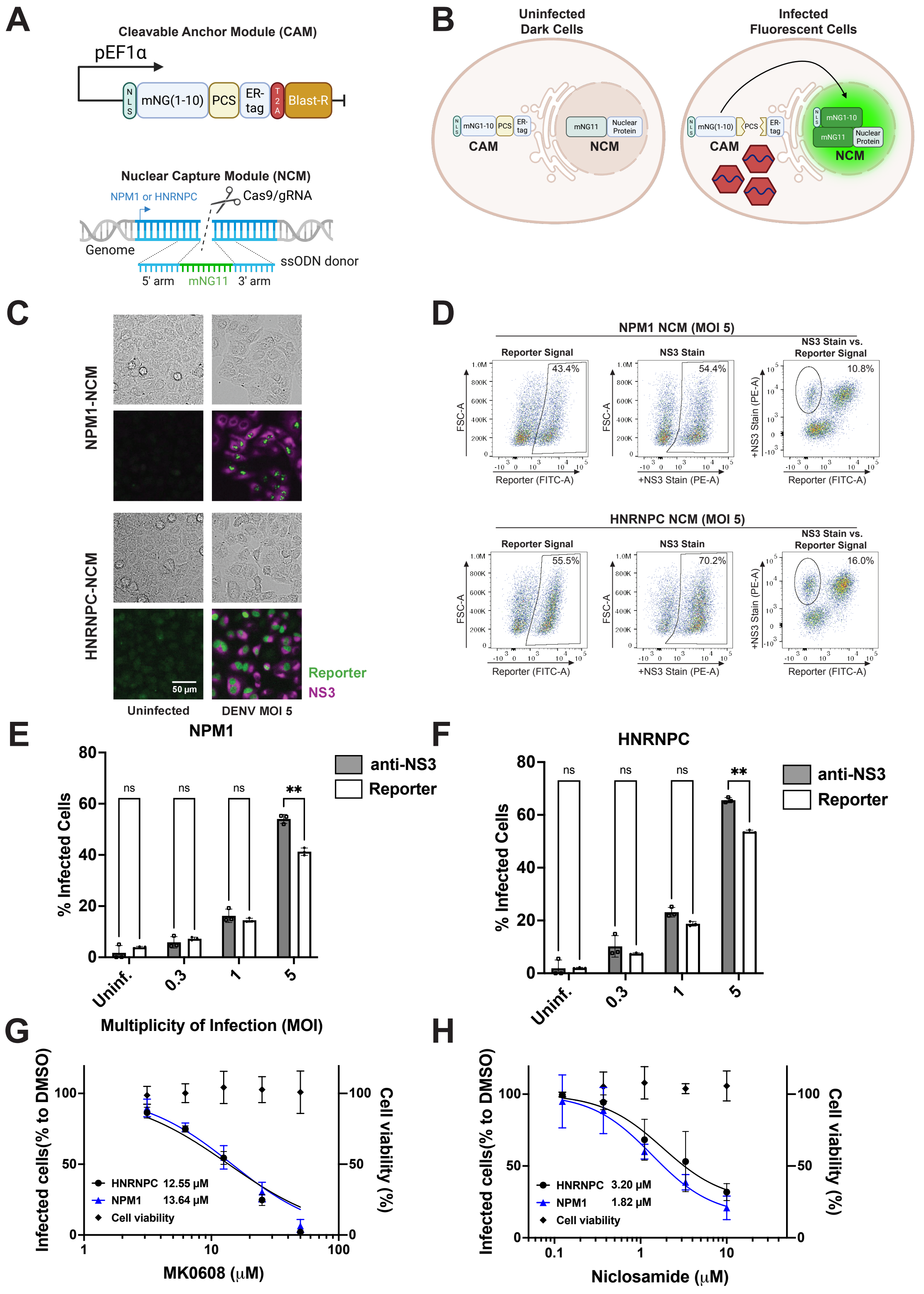

A
Cleavable Anchor Module (CAM)
pEF1α
mNG(1-10)
PCS
ER-tag
Blast-R
Nuclear Capture Module (NCM)
NPM1 or HNRNPC
Cas9/gRNA
Genome
ssODN donor
5' arm
mNG11
3' arm
B
Uninfected Dark Cells
Infected Fluorescent Cells
CAM
NCM
C
NPM1-NCM
HNRNPC-NCM
Reporter
NS3
50 µm
Uninfected
DENV MOI 5
D
NPM1 NCM (MOI 5)
Reporter Signal
NS3 Stain
NS3 Stain vs. Reporter Signal
43.4%
54.4%
10.8%
HNRNPC NCM (MOI 5)
55.5%
70.2%
16.0%
FSC-A
+NS3 Stain (PE-A)
Reporter (FITC-A)
E
NPM1
F
HNRNPC
anti-NS3
Reporter
% Infected Cells
Multiplicity of Infection (MOI)
Uninf.
0.3
1
5
G
Infected cells(% to DMSO)
Cell viability (%)
HNRNPC 12.55 µM
NPM1 13.64 µM
Cell viability
MK0608 (µM)
H
HNRNPC 3.20 µM
NPM1 1.82 µM
Niclosamide (µM)

**Supplementary Figure 1. Design and validation of a split-fluorescent reporter for DENV protease activity.**

(A–B) Architecture and activation of the two-component reporter. The Cleavable Anchor Module (CAM) contains NLS–mNG(1–10), a DENV protease cleavage site and an ER anchor. The Nuclear Capture Module (NCM) consists of endogenous NPM1 or HNRNPC tagged with mNG(11) by CRISPR knock-in. (C) Representative images of uninfected and DENV-infected A549 reporter cells (MOI 5), showing reporter fluorescence (green) and NS3 immunostaining (magenta). NPM1-NCM and HNRNPC-NCM cells are shown above and below, respectively. Scale bar, 50 µm. (D) Representative flow cytometry plots at MOI 5, showing reporter fluorescence, NS3 immunostaining and their relationship. Percentages of NS3-positive/mNG-negative cells are indicated. (E–F) Infection detected by NS3 immunostaining (grey) or reporter fluorescence (white) at the indicated MOIs in NPM1-NCM (E) and HNRNPC-NCM (F) cells. Each cell line was analyzed by two-way repeated-measures ANOVA with matching in both factors and Geisser–Greenhouse correction, followed by Šídák-adjusted comparisons between detection methods within each MOI. Exact P values are provided in Source Data. (G–H) MK0608 (G) and niclosamide (H) dose–response curves in both reporter cell lines. MK0608 absolute IC50 values were 13.64 µM (95% profile-likelihood CI, 11.78–15.77 µM) for NPM1-NCM and 12.55 µM (11.13–14.11 µM) for HNRNPC-NCM. Niclosamide values were 1.83 µM (1.11–3.03 µM) and 3.20 µM, respectively; the HNRNPC-NCM lower confidence limit was 1.63 µM, but the upper limit could not be determined. Infection is expressed relative to mean DMSO-control infection. Absolute IC50 is the fitted concentration giving 50% of this response. Viability is shown on the right axis as measured points without a fitted curve. Technical replicates were averaged within experiments before fitting infection-response curves as described in Methods. Quantitative data are mean ± s.d. from three independent experiments.

A

B

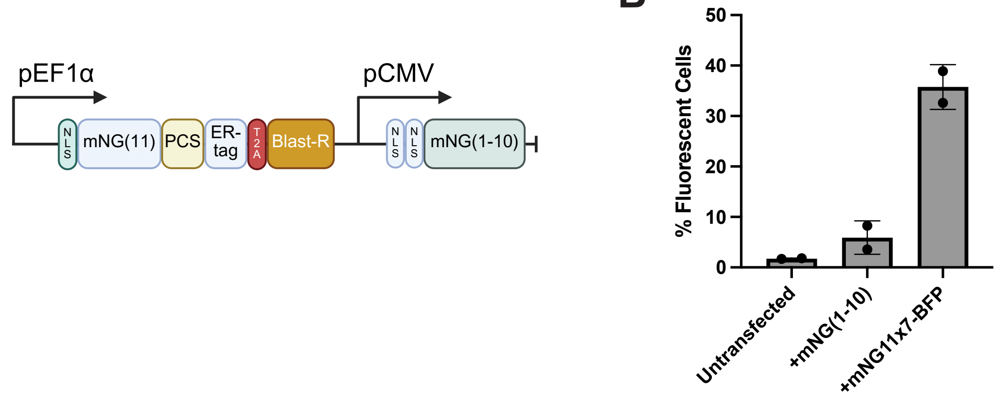
pEF1α
N L S
mNG(11)
PCS
ER-tag
T 2 A
Blast-R
pCMV
N L S
N L S
mNG(1-10)
50
40
30
20
10
0
% Fluorescent Cells
Untransfected
+mNG(1-10)
+mNG11x7-BFP

**Supplementary Figure 2. Stability of the split-mNG fragments in the initial single-vector reporter configuration.**

(A) Schematic of the initial single-vector reporter design in which mNG(1–10) was constitutively expressed in the nucleus (NCM) and the shorter mNG(11) peptide served as the ER-anchored cleavable element (CAM). (B) Fluorescence complementation assay assessing the stability of each split-mNG fragment in the initial reporter configuration. A549 cells expressing the initial single-vector reporter were transiently transfected with a plasmid encoding either mNG(1–10) or mNG(11)x7–BFP. Points show both independent biological experiments, with one technical measurement per condition per experiment; bars and error bars show mean ± s.d. (n = 2). No inferential statistical test was performed.

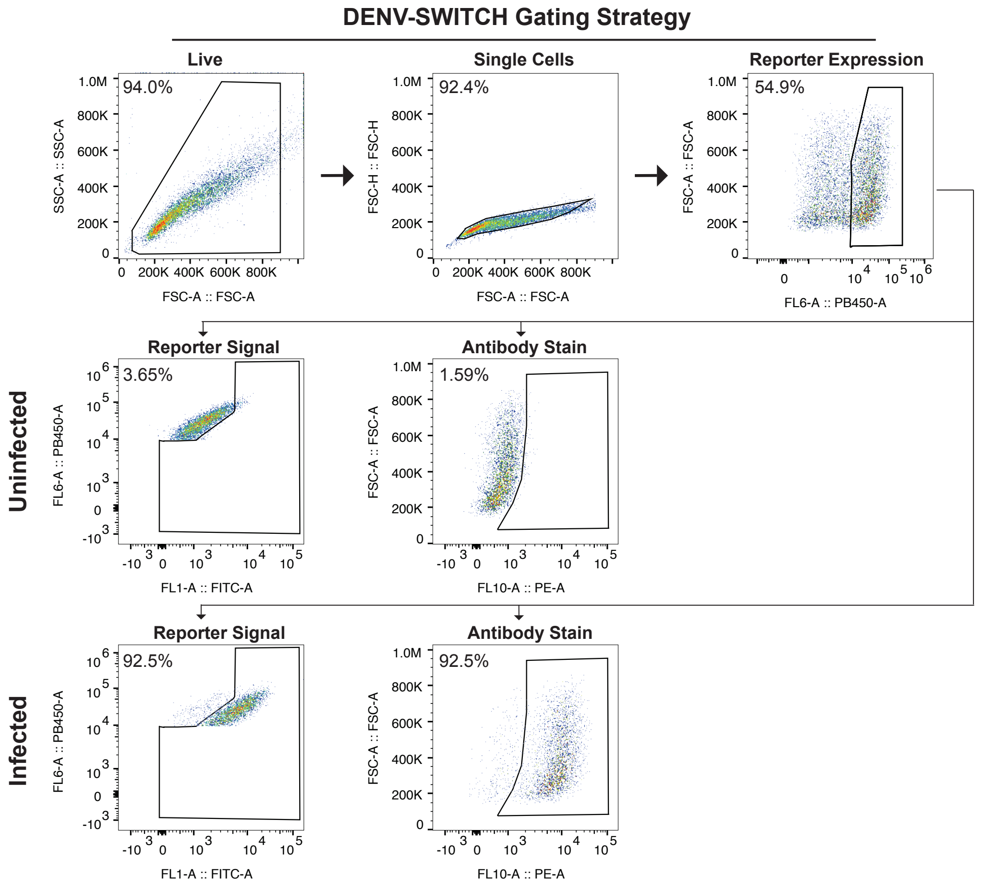
DENV-SWITCH Gating Strategy
Live
94.0%
SSC-A :: SSC-A
FSC-A :: FSC-A
Single Cells
92.4%
FSC-H :: FSC-H
FSC-A :: FSC-A
Reporter Expression
54.9%
FSC-A :: FSC-A
FL6-A :: PB450-A
Uninfected
Reporter Signal
3.65%
FL6-A :: PB450-A
FL1-A :: FITC-A
Antibody Stain
1.59%
FSC-A :: FSC-A
FL10-A :: PE-A
Infected
Reporter Signal
92.5%
FL6-A :: PB450-A
FL1-A :: FITC-A
Antibody Stain
92.5%
FSC-A :: FSC-A
FL10-A :: PE-A

**Supplementary Figure 3. Flow cytometry gating strategy for V-SWITCH reporter quantification.**

Representative flow cytometry plots from one experiment illustrating the sequential gating strategy used for V-SWITCH quantification in DENV-SWITCH A549 cells. Cells were first gated on forward scatter (FSC-A) versus side scatter (SSC-A) to exclude debris and dead cells, followed by FSC-A versus FSC-H to exclude doublets. BFP-positive cells defined the reporter-expressing population. Reporter-OFF and NS3-positive gates were set using uninfected samples in PB450-A (BFP) versus FITC-A (mNG) and PE-A (NS3) versus FSC-A plots, respectively. This panel illustrates the analysis procedure and was not statistically analyzed.

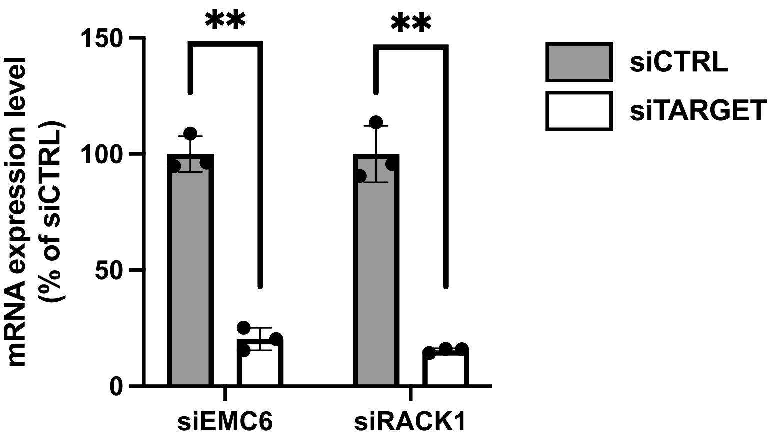

**
**
150
100
50
0
mRNA expression level
(% of siCTRL)
siEMC6
siRACK1
siCTRL
siTARGET

**Supplementary Figure 4. Validation of EMC6 and RACK1 silencing in A549 cells.**

RT–qPCR quantification of EMC6 and RACK1 mRNA levels in A549 cells transfected with siEMC6, siRACK1, or non-targeting control (siCTRL) for 72 h. Technical replicates were averaged before ΔCq calculation relative to GAPDH. Plotted expression was calculated as $2^{-\Delta Cq}$, normalized to the arithmetic mean of the corresponding siCTRL values, and expressed as a percentage. Inference was performed on paired ΔCq values using a paired two-tailed t-test for each prespecified gene-specific validation experiment; $P = 0.003364$ for EMC6 and $P = 0.001387$ for RACK1. No multiplicity correction was applied across these distinct gene-specific endpoints. Data are mean ± s.d. ($n = 3$ independent experiments).

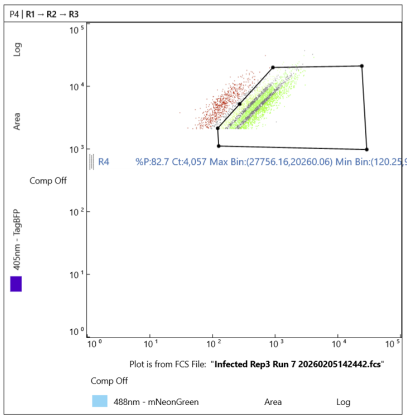
P4 | R1 → R2 → R3
405nm - TagBFP
Area
Log
Comp Off
R4
%P:82.7 Ct:4,057 Max Bin:(27756.16,20260.06) Min Bin:(120.25,951.73)
Plot is from FCS File: "Infected Rep3 Run 7 20260205142442.fcs"
Comp Off
488nm - mNeonGreen
Area
Log

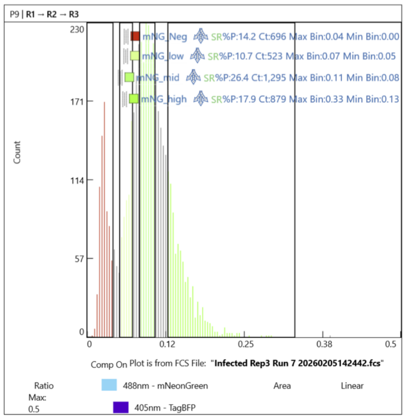
P9 | R1 → R2 → R3
Count
mNG_Neg SR%P:14.2 Ct:696 Max Bin:0.04 Min Bin:0.00
mNG_low SR%P:10.7 Ct:523 Max Bin:0.07 Min Bin:0.05
mNG_mid SR%P:26.4 Ct:1,295 Max Bin:0.11 Min Bin:0.08
mNG_high SR%P:17.9 Ct:879 Max Bin:0.33 Min Bin:0.13
Comp On
Plot is from FCS File: "Infected Rep3 Run 7 20260205142442.fcs"
Ratio
Max:
0.5
488nm - mNeonGreen
Area
Linear
405nm - TagBFP

**Supplementary Figure 5. FACS sorting strategy for RT-qPCR analysis of DENV RNA levels.**

Representative flow cytometry plots from one experiment showing the gating strategy used to sort A549 DENV-SWITCH cells into four populations based on the mNG/BFP fluorescence-intensity ratio: mNG-negative, low mNG, mid mNG, and high mNG. Cells were infected with DENV at MOI 5 and sorted at 24 hpi on a Bigfoot (ThermoFisher) cell sorter. Sorting was performed across three independent experiments, with 10,000 cells collected per gate. This panel illustrates the sorting procedure and was not statistically analyzed; the number of collected cells is not the biological sample size.

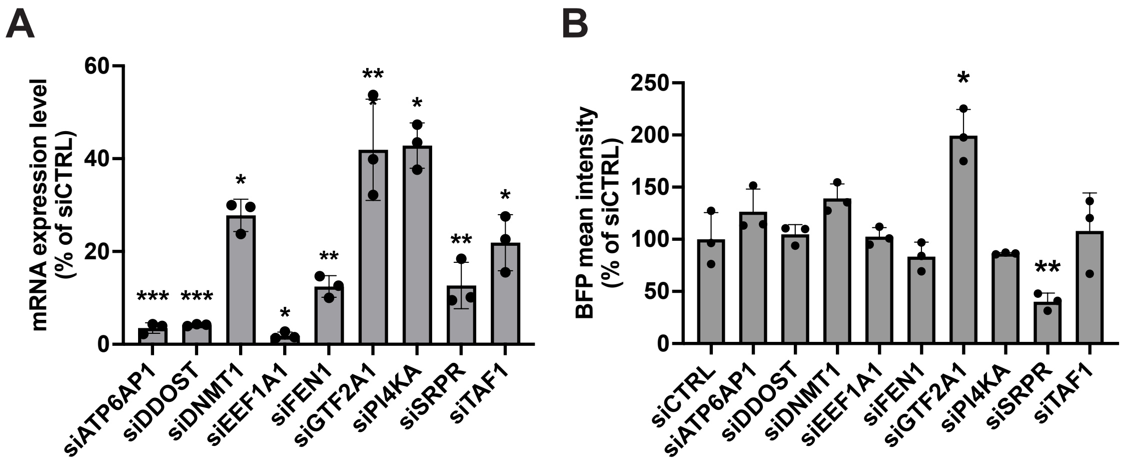
A
mRNA expression level (% of siCTRL)
0
20
40
60
***
***
*
*
**
**
*
**
*
siATP6AP1
siDDOST
siDNMT1
siEEF1A1
siFEN1
siGTF2A1
siPI4KA
siSRPR
siTAF1
B
BFP mean intensity (% of siCTRL)
0
50
100
150
200
250
*
**
siCTRL
siATP6AP1
siDDOST
siDNMT1
siEEF1A1
siFEN1
siGTF2A1
siPI4KA
siSRPR
siTAF1

**Supplementary Figure 6. Validation of the silencing efficiency of prioritized hits in A549-DENV-SWITCH cells.**

(A) RT–qPCR quantification of the prioritized hits' mRNA levels in A549 DENV-SWITCH cells transfected with the designated siRNA or non-targeting control (siCTRL) for 72 h. Technical replicates were averaged before ΔCq calculation relative to GAPDH. Plotted expression was calculated as $2^{-\Delta Cq}$, normalized to the arithmetic mean of the corresponding gene-specific siCTRL values, and expressed as a percentage. Inference was performed on paired ΔCq values using a paired two-tailed t-test for each prespecified gene-specific validation experiment; no multiplicity correction was applied across genes. P values were 0.000523, 0.000202, 0.010791, 0.018170, 0.005641, 0.006740, 0.026323, 0.002269, and 0.017980 for ATP6AP1, DDOST, DNMT1, EEF1A1, FEN1, GTF2A1, PI4KA, SRPR and TAF1, respectively. (B) Constitutive BFP reporter intensity after the same siRNA treatments. Inference was performed on log10-transformed raw fluorescence intensities using a Geisser–Greenhouse-corrected repeated-measures one-way ANOVA ($P = 0.0134$) followed by Dunnett's test versus siCTRL. Dunnett-adjusted P values for the targets in the order listed above were 0.2292, 0.9781, 0.1852, 0.9987, 0.2508, 0.0339, 0.9316, 0.0047, and 0.9890. Data are mean ± s.d. ($n = 3$ independent experiments).

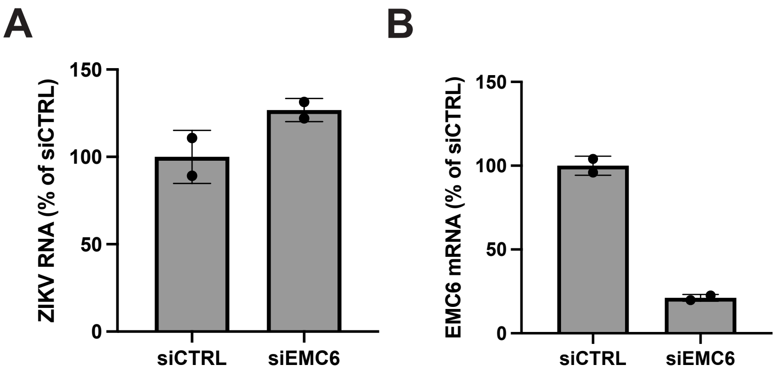
A
ZIKV RNA (% of siCTRL)
150
100
50
0
siCTRL
siEMC6
B
EMC6 mRNA (% of siCTRL)
150
100
50
0
siCTRL
siEMC6

**Supplementary Figure 7. Validation of EMC6 silencing effect on ZIKV infection in HeLa cells.**

(A) ZIKV RNA and (B) EMC6 mRNA levels measured by RT–qPCR in HeLa cells transfected with siEMC6 or non-targeting control (siCTRL) for 72 h, followed by ZIKV infection. Technical replicates were averaged before ΔCq calculation relative to GAPDH. Plotted values were calculated as $2^{-\Delta Cq}$, normalized to the arithmetic mean of the corresponding siCTRL values, and expressed as a percentage. ZIKV RNA was 126.8 ± 6.6% in siEMC6-treated cells versus 100.0 ± 15.2% in siCTRL cells; EMC6 mRNA was 21.2 ± 2.0% versus 100.0 ± 5.7%, respectively. Data are mean ± s.d. from two paired independent biological experiments. Owing to $n = 2$, no inferential statistical test is reported.

### Supplementary Tables and Methods

#### Supplementary Tables

**Supplementary Table 1. siRNA reagents used in this study.**

Target genes, suppliers, reagent identifiers and both strand sequences for the 11 gene-targeting siRNAs used for validation. Sequences are written 5′–3′; lowercase letters denote RNA and uppercase letters denote DNA. Strand 1 and Strand 2 correspond to the supplier's SEQ1 and SEQ2 labels. RACK1 is listed under its supplier identifier, GNB2L1. The Dharmacon ON-TARGETplus non-targeting control pool is identified by catalogue number; its component sequences were not supplied.

**Supplementary Table 2. Primer sequences used in this study.**

Primer names, applications and sequences for RT–qPCR, dial-out PCR, sgRNA-library amplification, indexing and sequencing. All sequences are written 5′–3′. The table includes the P5/P7 primer-pair assignments for the six CRISPR-screen samples and the custom Read 1 primer used to sequence the sgRNA.

**Supplementary Table 3. sgRNA sequences in the pooled CRISPRi library.**

Identifiers, target genes and 19-nucleotide spacer sequences for the 14,545 guides represented in the screen count matrix: 14,214 guides targeting 2,583 genes and 331 non-targeting controls. Sequences are written 5′–3′ with the original capitalization preserved. Guide identifiers and target labels match the count matrix; alternate annotations for identical spacer sequences are listed separately.

**Supplementary Table 4. Complete results of the pooled DENV CRISPRi screen.**

Gene-level results and raw sgRNA read counts for the Unsorted, Top and Low populations from two biological replicates. Gene-level results report Top-versus-Low beta coefficients, Wald Z scores, permutation and Wald P values and false discovery rates (FDRs), Figure 3B ranks and hit classifications. Negative beta coefficients and Z scores indicate depletion in Top relative to Low. Hits were defined by permutation FDR < 0.05; the nine hits with |Z| > 5 were selected for follow-up. ESRRG, SGK3 and the aggregated negative-control entry were excluded from estimation and are retained with explicit labels. Unsorted samples were retained for quality control but were not included in the Top-versus-Low model.

#### Supplementary Methods

##### Image-analysis quality checks and sensitivity analyses

**Complete-track analysis (Fig. 2B).** Cumulative reporter activation was calculated as 1 − S(t), with left truncation at each cell's first tracked frame and right censoring one frame before its final observation if activation was not detected. The processed tracking tables could not establish whether cells whose tracks ended early differed in their likelihood of subsequent reporter activation. We therefore repeated the analysis using only cells observed at every frame from 0 through 26. This assesses sensitivity to incomplete tracking but does not establish that tracking loss was independent of reporter activation.

**Sigmoid model and response groups (Fig. 2E–F).** The model was $R(t) = b + A/[1 + \exp(-k(t - t_0))]$, where b is baseline, A is rise amplitude, k is steepness and $t_0$ is the response midpoint. The fitted plateau was b + A and $t50 = t_0$. Frame-based midpoint estimates were converted to hours post-infection by adding 1.5 h. Cells with an observed transition to reporter positivity, a finite fitted plateau and source-fit $R^2 \geq 0.7$ formed the original amplitude-eligible cohorts: 1,656 cells in replicate 1 and 1,267 in replicate 2. Plateau-tertile cutoffs and the replicate-specific median plateaus used for normalization were calculated in these original cohorts and were not recalculated after timing-quality filtering. Plateau groups were not compared inferentially because grouping was defined by plateau itself.

**Coverage of the fitted response.** Timing-quality selection required at least eight observations, at least two measurements at or before fitted $t10 = t_0 - \ln(9)/k$, at least two at or after fitted $t90 = t_0 + \ln(9)/k$, and at least two measurements on each side of t50. Here, t10 and t90 denote the times at which the fitted response reaches 10% and 90% of its rise above baseline. The midpoint also had to lie at least one hour inside both observed track boundaries. Fits whose slopes reached the fitting bounds were excluded. These criteria require observations across the fitted rise but do not establish that a biological plateau was observed.

**Numerical fit checks.** To assess the reliability of the stored fits, diagnostic nonlinear least-squares refits used two starting points and the constraints $b \geq 0$, $A \geq 0$, $0.001 \leq k \leq 5$, with $t_0$ within the observed track. Refits had to agree with the stored solution: the fitted amplitude had to be positive, the relative reduction in residual sum of squares could not exceed $10^{-4}$, and the midpoint shift could not exceed 0.1 h. Fits with unresolved optimization or profile failures were excluded. These diagnostic refits were used for quality assessment and did not replace the stored estimates used in the figure.

**Midpoint uncertainty.** For each fixed value of $t_0$, b, A and k were reoptimized to construct an approximate 95% profile interval. The residual sum-of-squares cutoff was $SSEmin \times [1 + F_{0.95}(1, n - 4)/(n - 4)]$, where SSEmin is the minimum residual sum of squares and n is the number of observations. The primary analysis required a closed interval no wider than four hours; two- and six-hour limits were examined as sensitivity analyses. These intervals describe uncertainty within an individual fitted curve, assuming independent Gaussian residuals with constant variance. Correlation between successive measurements may make the intervals too narrow. They do not measure uncertainty across biological replicates, and midpoint and plateau estimates derived from the same curve are not statistically independent.

**Endpoint-fluorescence analysis.** As an exploratory comparison with the 24-hpi sorting experiment, we related t50 to the measured mNG/BFP ratio at 24.5 hpi (frame 23), with 23.5 hpi (frame 22) as an adjacent-frame sensitivity check. Endpoint-positive cells had to exceed the activation threshold at that time and have a confirmed two-frame activation run beginning at or before it. Endpoint-ratio tertiles were defined within each replicate before restricting cells by observed activation and timing quality, and these assignments were then held fixed. These groups were an imaging analogue of the sorting comparison, not the experimental FACS gates. At 24.5 hpi, 268 and 304 timing-quality cells were endpoint-positive in replicates 1 and 2. Their ratio–t50 associations were weakly negative (Spearman's $\rho = -0.045$ and $-0.147$) and sensitive to the quality-filtering rule.

**Midpoint-aligned fluorescence traces.** Observed time courses were aligned to t50 over −4 to +4 h. Linear interpolation was allowed only between observed measurements separated by no more than one hour; no extrapolation was used. Cells had to contribute at every aligned time point, retaining 280 cells in replicate 1 and 327 in replicate 2. The traces retained the measured response amplitudes and original plateau-group assignments. Because those groups were defined by amplitude, separation of the aligned traces was partly expected and was not treated as an independent test.
